# Cortex-wide Dynamics of Internal Decisions About Behavioral Context

**DOI:** 10.64898/2025.12.18.694943

**Authors:** Joshua Calder-Travis, Ruud L. van den Brink, Saanchi Thawani, Lars Schwabe, Tobias H. Donner

## Abstract

Many human decisions manifest in the outside world as motor actions. Correspondingly, neural signals reflecting decision formation are expressed in neural populations encoding the final action. Some decisions, however, are internal, shaping overt behavior indirectly through the selection of policies or rules that guide elementary sensory-motor decisions. We studied the human brain dynamics underlying internal decisions about the stimulus-response mapping rule for reporting visual orientation judgments. The rule changed unpredictably in a hidden fashion, and participants tracked it by accumulating ambiguous sensory cues. Computational model-based decoding of source-level magnetoencephalography data uncovered representations of the internal belief about the active rule in many areas, including occipital cortex. These representations exhibited profiles as predicted by normative models and as observed in action-related frontal areas during overt decisions. We conclude that a widely distributed network of cortical areas implements evidence accumulation for internal decisions, and that even visual cortex may represent high-level beliefs.

## Introduction

For intelligent behavior, agents must keep track of the current behavioral context to guide elementary sensory-motor decisions. In other words, inferences about the context are not directly mapped on specific actions, but instead shape elementary decisions, including decision policies and the mapping from specific features of the sensory input to specific motor outputs (Miller & Cohen, 2001). For example, the stimulus of a phone call from one’s boss may be mapped to picking up the phone in the context of the boss being in a good mood, but to rejecting the call in the context of the boss being in a bad mood.

Humans and other animals are capable of such context-dependent decision-making (Mante et al., 2013; Miller & Cohen, 2001; Sakai, 2008). This is true even when the context is changing in a hidden fashion, and decision-makers only receive ambiguous or noisy information about the context (Purcell & Kiani, 2016; Sarafyazd & Jazayeri, 2019; van den Brink et al., 2023). Such situations require agents to accumulate noisy evidence regarding the current context and to use this evidence for higher-order, internal decisions, such as the selection of the appropriate input-output mapping for an elementary decision (Purcell & Kiani, 2016; Sarafyazd & Jazayeri, 2019; van den Brink et al., 2023). The goal of the current study was to uncover how and where the beliefs supporting higher-order internal decisions are encoded in the brain.

Psychology and neuroscience have developed a good understanding of the computational and neural bases of elementary sensory-motor decisions under uncertainty based on sequences of noisy sensory input (Bogacz et al., 2006; Glaze et al., 2015; Gold & Shadlen, 2007; Murphy et al., 2021; Peixoto et al., 2021). Sequential sampling models, in which observers accumulate incoming sensory evidence for the possible responses over an extended period of time, describe an algorithm that enables them to negate the effects of partially unreliable evidence (Bogacz et al., 2006). These models have successfully fit behavior in a wide variety of tasks, suggesting that they are a good description of the underlying algorithm (Bogacz et al., 2006; Gold & Shadlen, 2007; Ratcliff & McKoon, 2008). This even includes challenging situations, in which the environmental state to be inferred by the decision-maker may change in a hidden manner (Filipowicz et al., 2020; Glaze et al., 2015; Murphy et al., 2021). In such situations, the normative algorithm (Glaze et al., 2015) also involves accumulation of evidence over time. But now, recent evidence needs to be weighted more heavily than older evidence, which may not be relevant any longer due to a recent state change; and evidence received during periods of high uncertainty or when there is a high probability that the state has just changed (also known as change-point probability, or CPP) needs to be dynamically up-weighted (Murphy et al., 2021). This latter property allows observers to rapidly switch belief, when previously accumulated evidence may be misleading.

Candidate neural decision variables (DVs) for elementary sensory-motor decisions involving evidence accumulation have been identified in populations of neurons that also encode the corresponding action plans (Donner et al., 2009; Gold & Shadlen, 2007; Hanks et al., 2015; Khilkevich et al., 2024; Murphy et al., 2021; Peixoto et al., 2021; Shadlen & Kiani, 2013; Steinemann et al., 2024; Wilming et al., 2020). Activity in these neural populations encodes the strength and the sign of the accumulated evidence, and predicts the specific behavioral choice. Importantly, these neural DVs also correspond to the activity patterns observed during the planning of the corresponding action when that movement was instructed (Murphy et al., 2021; Shadlen & Newsome, 2001). For example, when the task requires saccadic choice reports, neural DVs are found in saccade-selective neural populations of the oculomotor control network including the lateral intraparietal area and frontal eye fields (Gold & Shadlen, 2001, 2007; Kiani et al., 2014; Shadlen & Newsome, 2001). When the task requires manual choice reports, neural DVs are found in the premotor and primary motor cortex (Donner et al., 2009; Murphy et al., 2021; Peixoto et al., 2021; Wilming et al., 2020). This correspondence between neural DV and neural action plan is intuitive because, in such tasks, the evolving decision can be mapped directly on a motor action.

By contrast, little is currently known about where and how higher-order, internal decisions are processed in the brain despite such decisions being crucial for intelligent behavior, in particular for the context-dependent decision-making that humans and other primates can perform reliably (Mante et al., 2013; Miller & Cohen, 2001; van den Brink et al., 2023). Here, we explored the computational and neural basis of internal decisions about context, which could change in a hidden fashion and which governed the input-output mapping to be used for reporting a lower-order decision. In our task, information about the context was provided by noisy sensory evidence.

We used magnetoencephalography (MEG) source imaging (Baillet, 2017) of cortical population activity combined with atlas-based, multi-area decoding techniques (Wilming et al., 2020) to comprehensively map the cortical distribution of neural codes of belief states reflecting the normative DV. Previous fMRI work using an analogous task found that the belief about the context was reflected in distinct patterns of intrinsic correlations between population codes for stimulus and action in sensory and action-related areas, respectively (van den Brink et al., 2023). This fMRI study also provided preliminary evidence for the local encoding of the normative DV in some cortical areas including, surprisingly, early visual cortex (van den Brink et al., 2023). Therefore, one might expect to find the DV for the internal decision encoded in multiple cortical areas including visual cortex, possibly as a result of feedback from downstream areas (Wilming et al., 2020). Critically, however, the limited temporal resolution of fMRI does not disambiguate the neural responses to individual sensory evidence samples, and those of the more slowly evolving DV (i.e. belief state) that is being updated with each evidence sample. We intended to fill this gap with our current MEG-based spatio-temporal mapping approach.

Our results indicate that human participants computed the internal decision about context following an approximately normative belief updating scheme previously established for sensory-motor decisions (Glaze et al., 2015). A hallmark signature of this computation was the dynamic up-weighting of evidence occurring during states of high uncertainty about the context, or under a high probability of a change in the context (CPP; Murphy et al., 2021). Pupil-linked arousal tracked the latter two variables (uncertainty and CPP). The DV (belief state) governing the internal decision was encoded in the population dynamics of many areas including parietal, temporal, and visual cortex. The dynamics of this neural code around each new sample accurately tracked the belief updating assumed by our behavioral model. Change-point probability and uncertainty are reflected across more widely distributed cortical regions, consistent with input from brainstem neuromodulatory systems. Our results provide a first glimpse on the cortex-wide dynamics of neural DVs for internal decisions.

## Results

During MEG recordings of neural population activity, participants (N=19) performed a hierarchical decision-making task in a volatile context (Figure 1). The lower level of the task required participants to judge the orientation (vertical or horizontal) of large and full-contrast grating stimuli and report their orientation judgment by button presses with either the left or the right index finger, using one of two complementary stimulus-response mappings (i.e., context; Figure 1B). The higher level of the task entailed selecting the currently active context, which varied throughout the experiment (Figure 1A). The active context was signaled by visual cues, a sequence of small dots presented in the upper or in the lower visual hemifield (for symmetry, each “cue” was a pair of such dots on the left and right from fixation). In one condition (“Instructed”), cues were unambiguous, occurring either in the lower visual hemifield for Context 1, or in the upper visual hemifield for Context 2. In another condition (“Inferred”), cues were ambiguous, with vertical positions drawn from two overlapping distributions (see Methods for details). In both conditions, the context could switch unpredictably, with a probability (hazard rate *H*) of 0.08 from one cue to the next. Determining the active context based on the cues was easy in the Instructed condition, but in the Inferred condition, this required sophisticated inference of the sort studied in previous work on lower-order sensory-motor decisions (Filipowicz et al., 2020; Murphy et al., 2021; Tavoni et al., 2022; Weiss et al., 2021). The Inferred condition required accumulating the noisy information provided by the cues in an adaptive fashion that strikes a balance between stable accumulation and sensitivity to change (Glaze et al., 2015).

**Figure 1:**
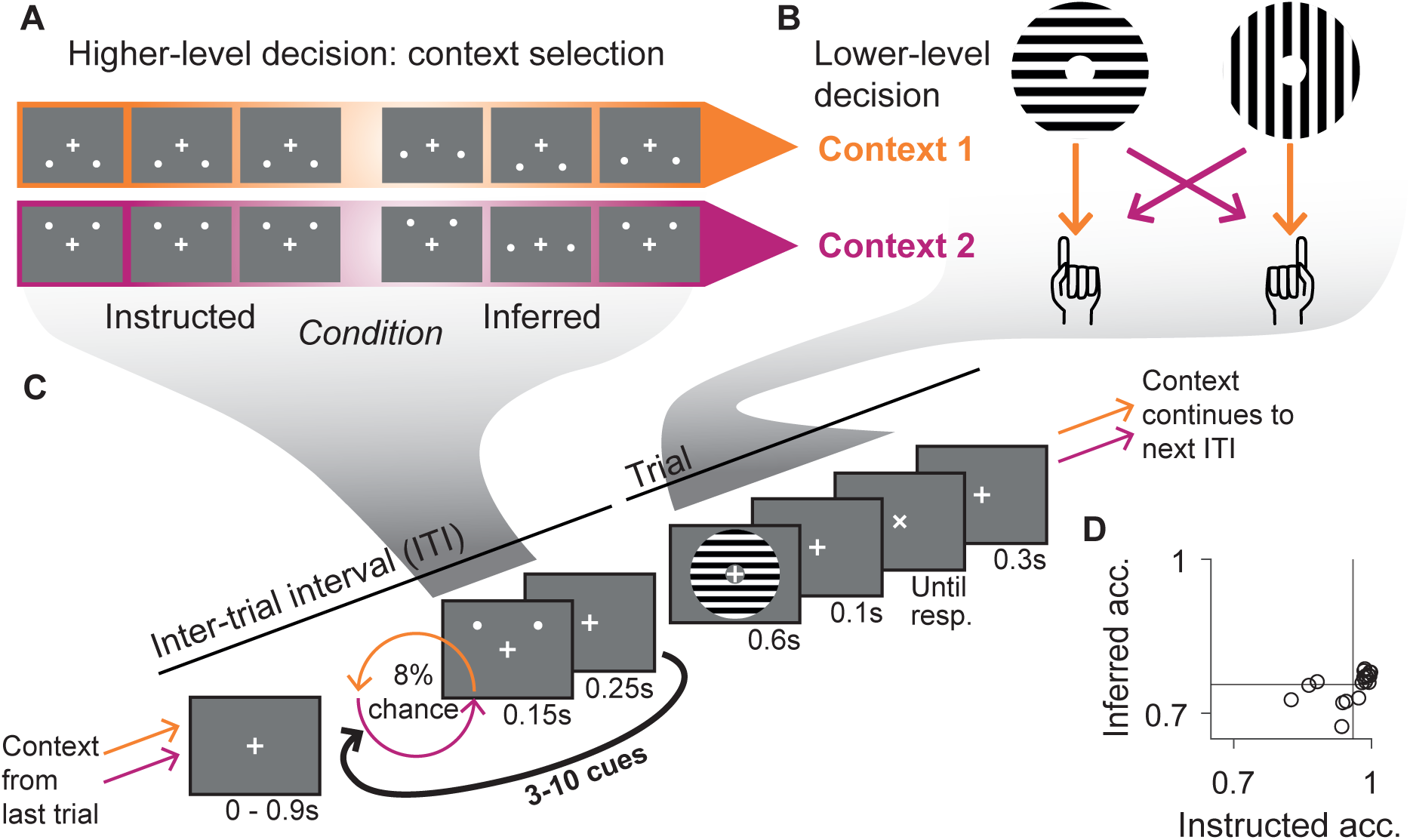
Hierarchical decision-making task. The task required decisions at two levels. (A) The higher-level task required selecting the appropriate stimulus-response mapping for the lower-level task, which flipped between two contexts. Visual cues – pairs of dots – below the horizontal midline were indicative of Context 1, and cues above the midline were indicative of Context 2. In the Instructed condition the cues were completely reliable. In the Inferred condition noise was added to the position of the cues. (B) On the lower-level, participants used the selected stimulus-to-response mapping to report a simple orientation judgment of a visual grating stimulus. (C) Combined sequence of task events consisting of an ITI during which participants were presented with between 3 and 10 cues, and a trial period, containing the response to the basic mapping task. Before each cue the context could potentially change. The context active at the end of one trial was the context active at the beginning of the next ITI. (D) Accuracy of participants in Instructed and Inferred conditions. Vertical and horizontal lines represent the mean accuracy across participants in the respective condition.

As expected, participants performed well above chance in both conditions, but consistently worse in the Inferred condition (*t*(18) = 20.1, *p* = 9.17 × 10^−14^, two-tailed, *d* = 4.60) reflecting the uncertainty inherent in this task (Figure 1D). This taps into the internal decision process that we dissect below, at the behavioral, computational, and neural levels.

### Participants accumulate noisy cues in an adaptive fashion

The normative algorithm for solving this task in the Inferred condition entails sequential updating of a belief about the active context with each sample of evidence (i.e., each cue), whereby the latter quantities are expressed as log-likelihood ratios for one over the other context (Glaze et al., 2015; Figure 2A). In each updating step, the prior belief (i.e., log-prior ratio) is combined with the evidence (log-likelihood ratio, LLR) into a posterior belief (log-posterior ratio) using Bayes rule. The posterior belief is then passed through a non-linear transform to become the prior belief for the next updating step, and so on, translating into a gradual accumulation of evidence. The shape of the non-linearity depends on the hazard rate *H*, which renders this updating scheme adaptive (Glaze et al., 2015). The model assumes that the internal decision about context at any moment is based on the sign of the log-posterior ratio. In other words, the belief state equals the normative decision variable for the higher-level decision.

**Figure 2:**
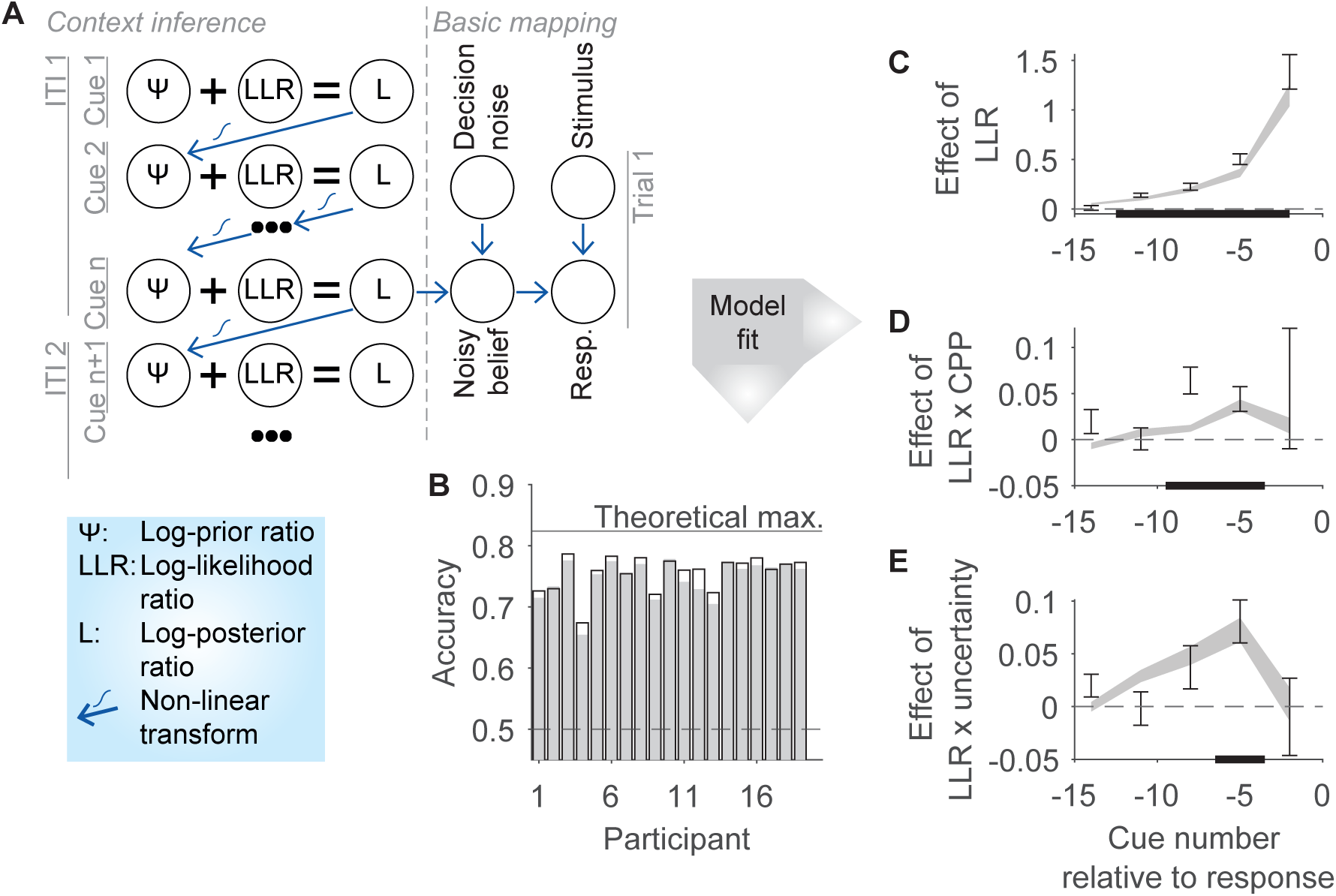
Modeling the internal context selection process in the Inferred condition. (A) Schematic of the normative (Bayes-optimal) solution for the higher-level decision. The algorithm involves combining a prior belief with incoming evidence (in the form of a log-likelihood ratio) and non-linearly transforming the resulting posterior to form the prior before the next cue (Glaze et al., 2015). During trials, a noisy version of the posterior belief is used to determine the current context, and hence, how to map the presented stimulus to a response. (B) Individual performance in the Inferred condition in the data (hollow bars) and in the model fits for a variant of the normative model that fit the data well (shading; “Miscalibrated h and b normative”), along with the theoretical maximum performance of a noiseless, calibrated and normative observer (see main text). (C-E) Temporal profiles of evidence weighting for internal decision (context selection), based on logistic regression (Inferred condition; see Methods). (C) Main effect of LLR on internal decision. (D, E) The regression also included terms to assess the modulatory effect of change point probability (CPP) and uncertainty. Error bars, participant data; gray shading, model fits (“Miscalibrated h and b normative”). (C-E) Error bar and shading widths correspond to *±*1 standard error of the mean across participants. Bold lines on x-axis, significant difference from zero (*p <*= 0.05; two-sided) of participant data points in a permutation cluster-based T-test with threshold-free cluster enhancement (see Methods).

The accuracy of all participants in the Inferred condition was lower than the theoretical maximum predicted by the exact normative algorithm described above (Figure 2B), implying that their performance was limited by suboptimalities. Even so, three lines of evidence indicate that participants used the normative strategy illustrated in Figure 2A, subject to noise in the transformation from belief to context selection, idiosyncratic biases in the representation of the context volatility (i.e., *H*) and potentially also in the mapping from cue positions to LLRs (*β*). First, the normative model with the above suboptimalities accurately captured most participants’ performance (Figure 2B; “Miscalibrated h and b normative” model in Methods). Non-adaptive strategies (i.e., strategies ignoring context volatility) such as basing the context selection only on the last sample or perfect integration (i.e., accumulating the entire evidence sequence with equal weight) yielded substantially worse performance incompatible with the data (Figure 2 – Supplemental Figure 1A; Glaze et al., 2015). Two simpler adaptive strategies, a leaky linear accumulation with adaptive leak (Glaze et al., 2015; Ossmy et al., 2013) and in particular perfect integration with non-absorbing bound (Glaze et al., 2015) performed more closely to the data (Figure 2 – Supplemental Figure 1A), which is expected because those two strategies (especially the latter, for our task parameters) approximate the normative strategies (Glaze et al., 2015). Formal model comparisons favored variants of the normative strategy (although the model with non-absorbing bound did not perform significantly worse; see “Bounded accu.” in Figure 2 – Supplemental Figure 1B, 1D; all compared models featured noise in the transformation from belief to context selection). Unless otherwise noted, we focus on and use the variant of the normative model with potentially miscalibrated hazard rate (*H*), and mapping from cue positions to LLRs (*β*), which fit best according to the AIC (Figure 2 – Supplemental Figure 1B; see Methods).

Second, the temporal profiles of the weighting of evidence samples (i.e. cues) in the internal decision (i.e., context selection) exhibited key signatures of non-linear, dynamic evidence upweighting, which previous work has identified as diagnostic for the normative accumulation scheme (Murphy et al., 2021; Figure 2C-E): Participants, and the fitted normative model, gave stronger weight to cues later than to cues earlier in the sequence (Figure 2C); they also gave stronger weight to cues occurring during states of high uncertainty (i.e. log-prior belief close to 0; Figure 2E) or when the change-point probability (CPP) estimated from the normative model with true (not fitted) parameters was high (Figure 2D; Methods). All three signatures were accurately captured by the fitted normative model (Figure 2C-E; gray shading), and together are inconsistent with all alternatives discussed above (including accumulation with adaptive leak) except accumulation to a non-absorbing bound (Murphy et al., 2021).

Third, also replicating previous work on elementary decisions (Murphy et al., 2021, 2024), we found that participants’ pupil responses to individual cues were modulated by uncertainty and CPP gauged from the fitted normative model (Figure 3). Taken together, these observations support the notion that participants used an adaptive computation that, while subject to biases and noise, approximated the normative strategy for the task at hand. We, therefore, used the normative model fits to unravel representations of task-relevant latent variables in cortical population activity in the following.

**Figure 3:**
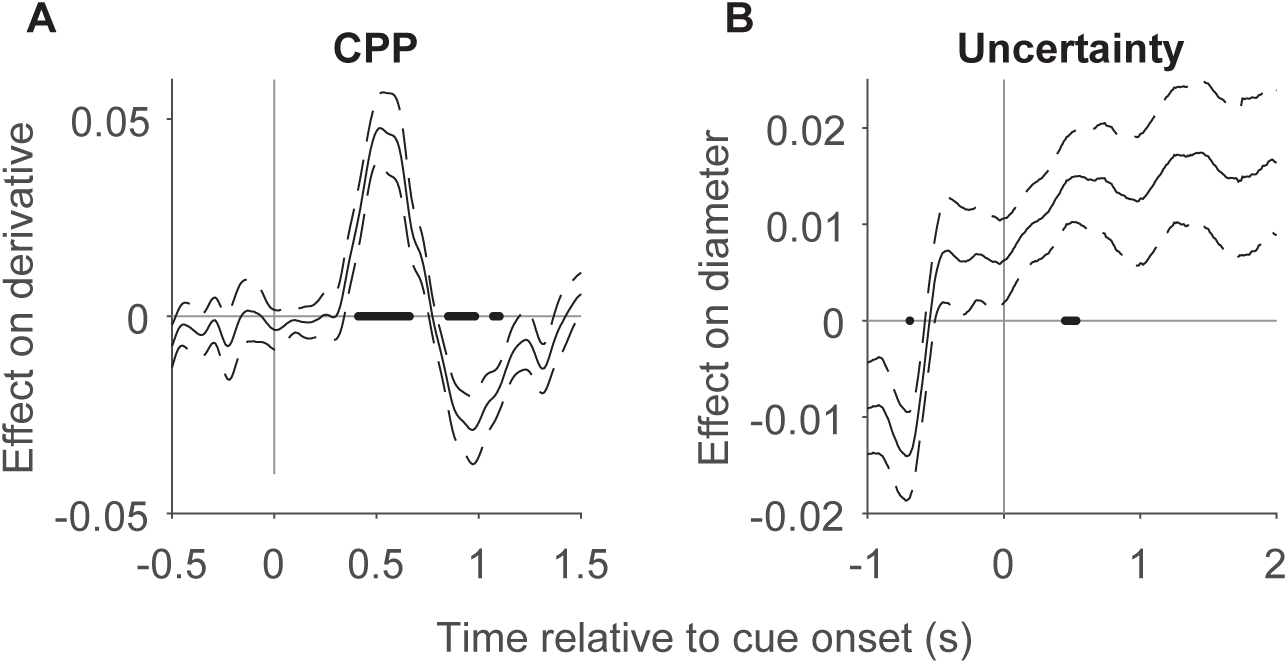
Latent behavioral model variables modulate pupil responses to individual cues. (A) The effect of CPP on the derivative of the pupil diameter time course (preregistered), and (B) the effect of uncertainty on the pupil diameter time course, both time-locked to cue onset (Inferred condition) and quantified in terms of regression coefficients (Methods). Dashed lines, *±*1 standard error of the mean. Bold lines on the axis, intervals in which the data were significantly different from zero. Significant differences from zero assessed in (A) with non-parametric permutation test, with false discovery rate correction (two-sided, *q <*= 0.05). No points in (B) were significant using this preregistered test. Significance in (B) corresponds to an alternative test against zero that accounts for the temporal structure of the data: A permutation cluster-based t-test with threshold-free cluster enhancement (*p <*= 0.05; two-sided; see Methods).

### Distributed coding of decision variable for internal decision

We applied a regionally-specific MEG decoding approach developed in previous work (Murphy et al., 2021; Wilming et al., 2020; Figure 4). We used MEG source imaging (Baillet, 2017) and an established anatomical parcellation (Glasser et al., 2016) to estimate spatial patterns of neural activity at each time point, within each of a number of regions covering the complete cortical surface, whereby we always combined the activity patterns from both homotopic regions of each hemisphere (results always shown on left cortical surface for visualization). For all main analyses we used a coarser parcellation comprising 22 area ‘groups’ per hemisphere (see Monov et al., 2024), but we verified in control analyses that the pattern of results also holds for the full, finer-grained parcellation (180 areas per hemisphere) described by Glasser et al. (2016). Following dimensionality reduction, these spatial patterns were used as features for cross-validated, multivariate decoding of the prior and the evidence (Methods). “Leave one-session out” cross-validation provided a very stringent test: Decoders were trained and evaluated on data collected from the same participant but on different days. Prior and evidence were estimated using the fitted normative model for each participant. We focused on three equally long intervals surrounding the presentation of each cue, to quantify the cortical distribution of the latent variables involved in each belief updating step (Figure 4A, B): just before cue presentation up to 25 ms following cue onset, so preceding any sensory cortical response (’pre-cue’), just after cue presentation (’early post-cue’) and following the removal of the cue from the display but preceding the presentation of the next cue (’late post-cue’).

**Figure 4:**
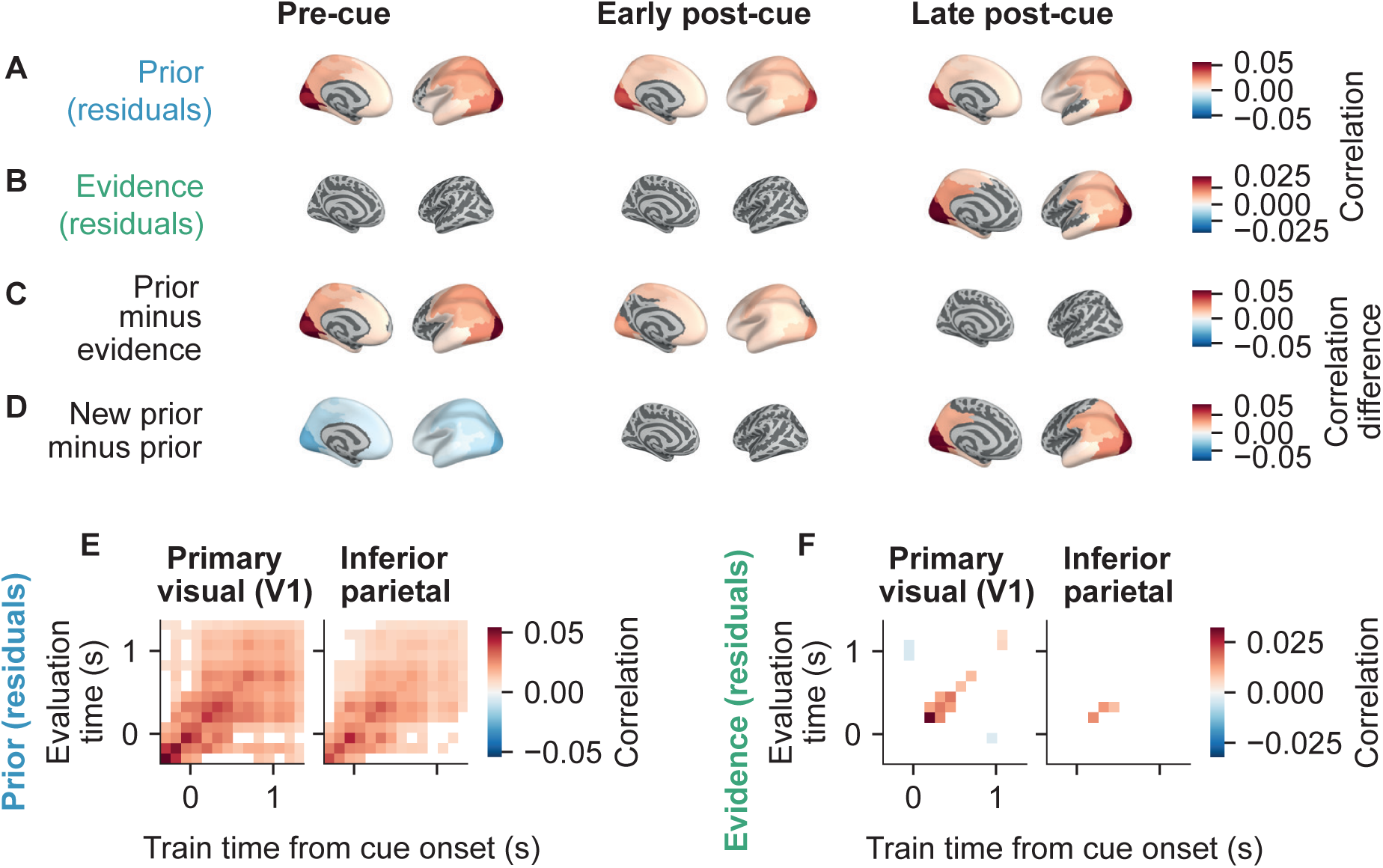
Decoding of prior belief about the context, and incoming evidence regarding the context, relative to the onset of cues in the Inferred condition. Decoding results are plotted separately for 22 cortical regions (homotopic regions from both hemispheres combined), and were produced using source activity estimates (Methods). Cross-validated performance was evaluated through the correlation between decoder output and the values of the target variables (prior or evidence). Regional decoding performance is shown on a reconstruction of the left cortical surface for visualization; decoding is always based on activity patterns from both hemispheres. (A) Decoding of log-prior ratio with effects of log-likelihood ratio (LLR) regressed out. (B) Decoding of evidence (i.e. LLR) with effects of log-prior ratio regressed out. (C) The difference in decoding performance for prior (residuals) and decoding performance for evidence (residuals). That is, the difference between (A) and (B). (D) The difference in decoding performance between the log-prior ratio after being updated with the current cue, and the log-prior ratio before being updated with the current cue. Positive values indicate that the updated prior is more strongly decoded than the old prior. (E, F) Temporal decoding generalization matrices for prior (residuals) and evidence (residuals) in the primary visual and inferior parietal regions, showing the cross-validated performance of decoders trained on data from a specific time point, evaluated on data from all other time points. In all panels, only regions (A-D) or time points (E, F) with decoding significantly different from zero are shaded (assessed using two-sided one-sample T-tests corrected using false discovery rate; *q <*= 0.05; see Methods).

Across all three intervals (and with little change between them), the prior was robustly decodable from a large number of areas spanning all four lobes, but centered on visual, parietal, and temporal cortices (Figure 4A). Notably, decoding of the prior during the pre-cue interval was significant even for the primary visual cortex (area V1). To further characterize the nature of the underlying neural codes, we focused on two regions of interest: Primary visual cortex V1, which must be involved in processing the visual-spatial cues because it is the entry point of visual information to cortical processing and contains a precise topographic map of the visual field (Wandell et al., 2007); and inferior parietal cortex, an area of association cortex that does not (different from areas of the intraparietal sulcus; Silver and Kastner, 2009) contain visual field maps. For both regions, temporal generalization analysis (King & Dehaene, 2014) showed that the code was largely stable throughout the time preceding and following the cue presentation (Figure 4E). By contrast, the evidence provided by the cue was encoded in a more transient code, from about 200 ms to about 700 ms following cue onset, and more robustly in V1 than inferior parietal cortex (Figure 4F). Correspondingly, we found significant evidence decoding for the late post-cue interval, peaking in visual cortical areas (Figure 4B).

Similar maps were obtained when running the analysis on a more fine-grained parcellation of cortical areas (180 areas per hemisphere; Figure 4 - Supplemental Figure 1) or when comparing the decoding performance in each region against the decoding performance obtained for primary auditory cortex (area A1; Figure 4 - Supplemental Figure 2).

Given the visual-spatial format of the evidence samples, one concern is that the prominent decoding of prior belief from activity in early visual cortical areas could be due to gaze shifts of participants tracking the cues with their eyes. We found small deviations of gaze positions from central fixation between intervals in which one or the other context was active (Figure 4 - Supplemental Figure 3A). While effects of eye movements were removed from the MEG signals using independent components analysis during preprocessing (Methods), this removal may be imperfect and residual effects of gaze position, or the resulting retinal shift of the fixation cross, could cause distinct patterns of neural responses in cortical areas with topographic visual field maps. Such a response would, in turn, correlate with prior belief and thus be picked up by our belief decoder. This scenario predicts a flip in the correlation between actual prior and the prior predicted by neural decoders, when the gaze position is inconsistent with the prior, a prediction refuted by the data (Figure 4 – Supplemental Figure 3). Specifically, we performed a median split of y-gaze position (average vertical position during the pre-cue time window) and a median split of the prior residuals. “Gaze consistent” refers to cues for which y-gaze position and the (residual) prior were on consistent sides of the median split. For example, gaze was in the upper half of the visual field, and prior was on the side of the median split that was associated with cues in the upper-half of the visual field. “Gaze inconsistent” refers to cues for which y-gaze position and the (residual) prior were on opposite sides of the median splits. If the decoders learn to decode gaze position, then performance on the gaze inconsistent trials should be negative: on these trials the gaze position is always misleading. Instead the pattern of results was highly similar across the two types of cue (Figure 4 – Supplemental Figure 3B and 3C). Indeed, the correlation of group average maps of decoding performance across areas (fine-grained parcellation using 180 parcels; all parcels included, whether or not significant) when evaluated on gaze consistent cues, with the average decoding performance maps on gaze inconsistent cues, was 0.85 in the pre-cue window (0.58 in the early post-cue window, and 0.66 for late post-cue window). These control analyses strongly argue against eye movements as a significant confound in the decoding of the internal decision variable.

The maps of prior and evidence decoding differed mainly during the pre-cue and early post-cue intervals, while they largely resembled each other in the late post-cue interval (Figure 4A,B), a visual impression that was confirmed by direct statistical comparison of both maps (Figure 4C). These results suggest the idea that the parietal, temporal, and visual cortical areas are involved in the complete belief updating computation entailing the representation of the prior belief and the new evidence sample, followed by the representation of the updated (posterior) belief resulting from the combination of prior and evidence described by the behavioral model. We used two approaches to test this idea. In the first approach, we subtracted the maps for decoding of the current prior from the maps for decoding of the next (i.e., updated) prior. Negative values on the maps during the pre-cue interval indicate that the current prior was more strongly decodable than the next prior, which cannot yet have been computed. Once the cue is presented then the next, updated, prior can be computed and this then becomes more strongly decodable, as indicated by the positive values on the maps. In line with the above idea that the complete belief updating computation occurs in parietal, temporal, and visual cortical areas, the maps are of similar topography (but opposite sign) for the pre-cue and late post-cue intervals (Figure 4D).

Our second approach was based on the insight that the adaptive (non-linear) belief updating entailed in the normative model can be approximated by the linear combination of the prior, the evidence, the evidence multiplied by the associated uncertainty, and the evidence multiplied by the associated CPP (Murphy et al., 2021). This linear decomposition of the belief updating entails the upweighting of evidence by associated uncertainty and CPP that we found to shape the internal decision behavior of our participants (Figure 2C-E). We, therefore, asked whether the areas in which activity reflected the updating of prior beliefs in the previous analysis, also exhibited a modulation of the neural representation of evidence samples (LLR) by the associated CPP and uncertainty. We found this to be the case, with spatial maps for the modulation of evidence (LLR) by CPP and uncertainty resembling those from our main analysis for the prior (compare Figure 4 - Supplemental Figure 4A and B with Figure 4A). The spatial maps for decoding these modulation terms – that may be viewed as components of the belief update – especially resembled the spatial map for regions with stronger decoding of the updated vs. previous prior, once the updated prior began to be more strongly decodable during the late post-cue period (compare Figure 4 - Supplemental Figure 4A and B with Figure 4D).

Taken together, patterns of neural population activity in visual and parietal cortex exhibited hallmark signatures of the updating of the decision variable (belief) postulated by the normative model for the higher-level decision in our task.

### Decoding of the decision variable in the absence of new visual evidence

The previous analyses used the intervals surrounding the onset of cues. At these times participants must represent their evolving belief while at the same time evidence is being presented on the display and must be processed by participants. An interesting comparison is the period of time leading up to responses, which follow presentation of the grating stimulus (Figure 1). Shortly before the response, the belief needs to be represented, as it guides the higher-order decision about the context (active rule), that is in turn used to report the lower-order decision with a button press (Figure 2). However, at the time of response there is no need to represent incoming evidence for the higher-order decision. Indeed no such evidence is presented for a period of at least 0.95s prior to a response (Figure 1).

For these reasons, we looked at the decoding of evidence and belief time-locked to responses instead of cues (Figure 5). For decoding of belief we decoded the posterior, instead of the prior, because it is the posterior that participants must use at this time. In Figure 5 the “late pre-response” window is a time window that immediately precedes the response. The posterior was significantly decodable from areas similar to those that had shown strong decoding of prior when time-locking to cues (Figure 5A). (Note that due to the task design there are far more cues than responses, meaning response-locked analyses will have reduced power to detect effects.) As expected, during this time-window there was no significant decoding of the evidence provided by the most recently presented cue (Figure 5B). The decoding of the posterior was significantly better than decoding of the evidence (Figure 5C). These results suggest a similar representation of belief, even during periods where new visual information is not being processed.

**Figure 5:**
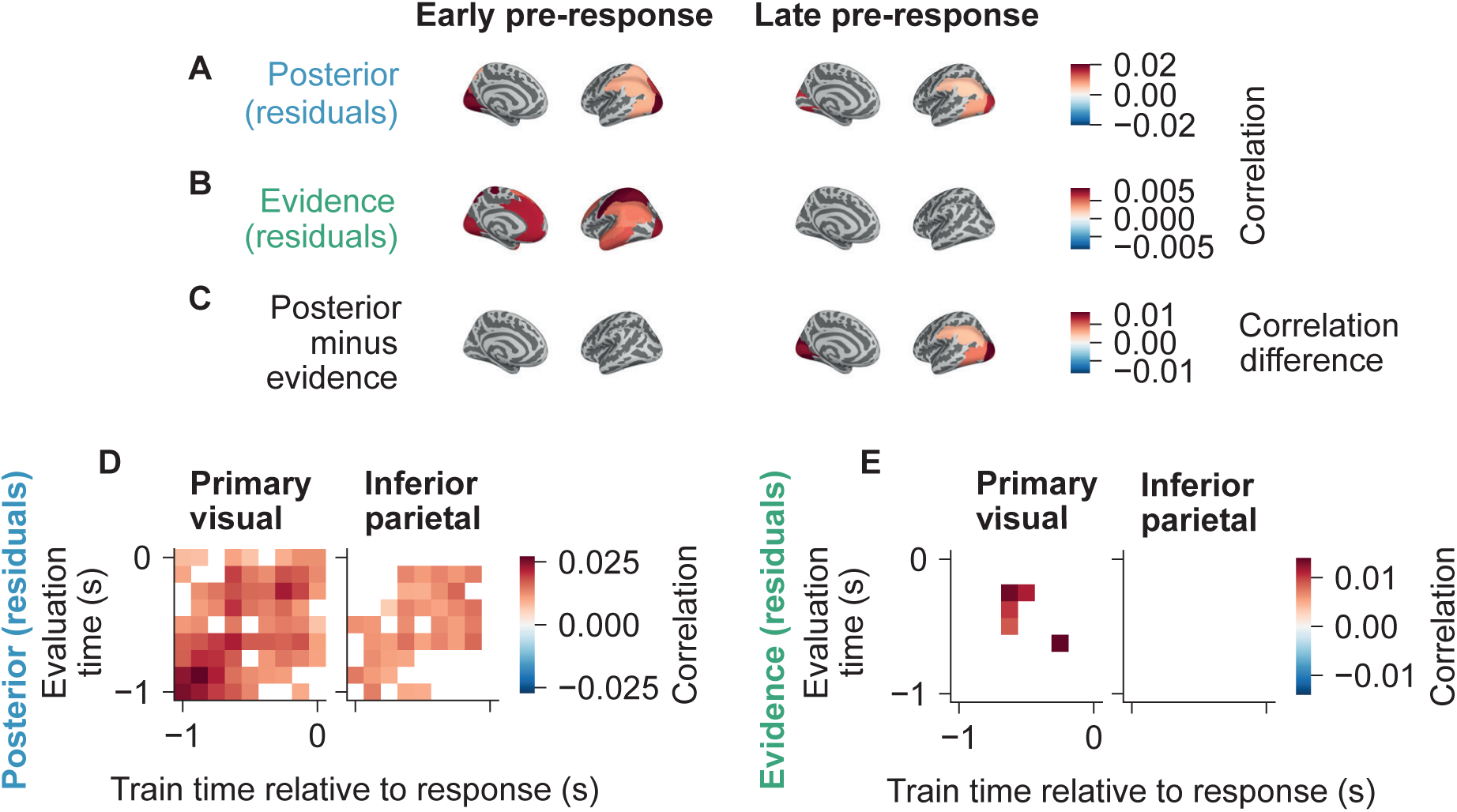
Decoding of prior belief about the context, and incoming evidence regarding the context, relative to responses (i.e. button presses) in the Inferred condition. Decoding was performed and evaluated analogously to that in Figure 4. (A – C) Decoding performance was evaluated through correlation. Only regions where decoding was significantly different from zero are shaded. (A) Decoding of log-posterior ratio (LPR) with effects of log-likelihood ratio (LLR) regressed out. (B) Decoding of evidence (i.e. LLR) with effects of LPR regressed out. (C) The difference in decoding performance for posterior (residuals) and evidence (residuals). That is, the difference between (A) and (B). (D and E) Decoding generalization matrices for posterior (residuals) and evidence (residuals) in the primary visual and inferior parietal regions. Only regions or time points with decoding significantly different from zero are shaded (assessed using two-sided one-sample T-tests corrected using false discovery rate; *q <*= 0.05). Further details in Methods.

Further adding to this picture are temporal generalization analyses (Figure 5D, E) which suggest that the representation of the posterior was stable over time, particularly in primary visual cortex (Figure 5D).

### Internal decision variable is expressed in alpha-band activity

Previous MEG work on elementary sensory-motor decisions has found representations of evolving decision variables in visual cortical field maps including V1, specifically expressed in the alpha (8-12 Hz) frequency band (Murphy et al., 2021). To test if the same held for the early visual cortical representation of the prior in our hierarchical task, we repeated the above decoding analyses on spatial patterns of spectrally resolved power estimates for each cortical area (see Methods). This showed little decoding precision in a very low-frequency band (5 *±* 2.5 Hz), but robust decoding of the prior in the alpha-band (10 *±* 2.5 Hz; Figure 6). (We estimated power at two individual frequencies, 5Hz and 10 Hz, but due to frequency smoothing, the estimates reflect activity at a range of frequencies.) This result highlights a further similarity in the nature of the DV representation for higher-order internal decisions, as studied here, and the DV representation for elementary sensory-motor decisions, as studied previously.

**Figure 6:**
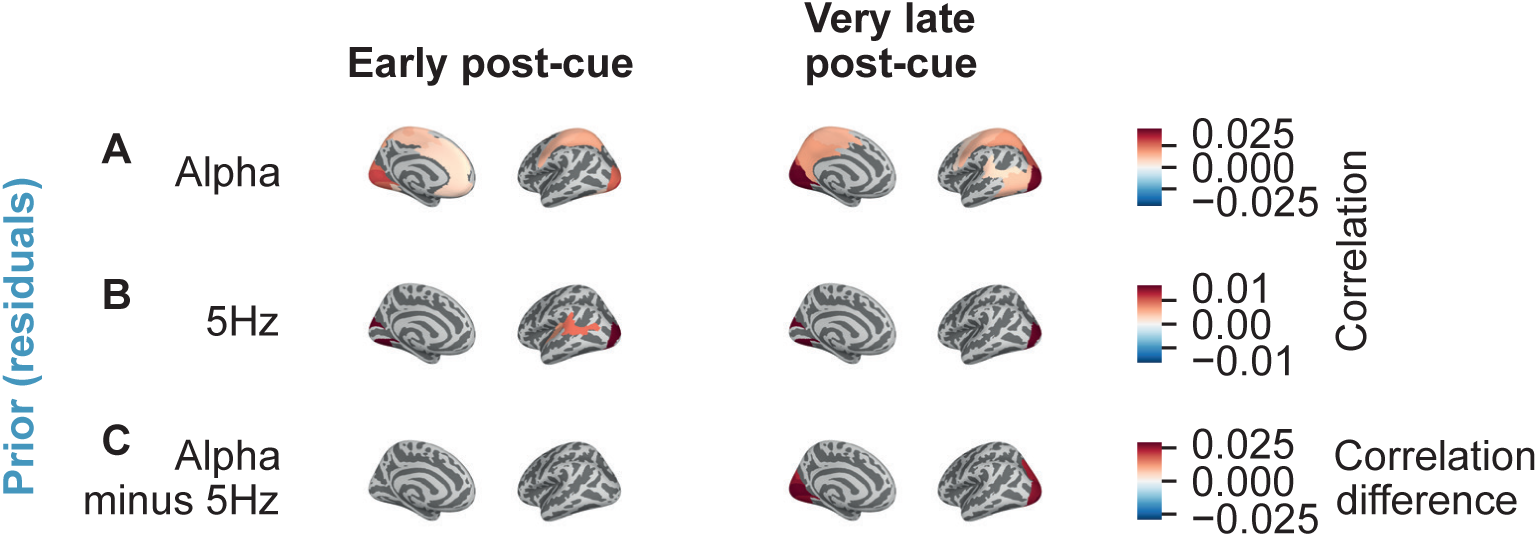
Decoding of prior in different frequency bands. Performed relative to the onset of cues in the Inferred condition. Decoding was performed separately for different brain regions, now using source estimates of spectral power in two frequency bands (5 *±* 2.5 Hz and 10 *±* 2.5 Hz) instead of the broadband time-domain signal. Otherwise decoder training and evaluation was analogous to that used in Figure 4. The target of the decoding was log-prior ratio with effects of log-likelihood ratio (LLR) regressed out. (A) Decoding using alpha power (10 Hz). (B) Decoding using power at 5 Hz. (C) The difference in decoding performance when using alpha power and when using 5 Hz. That is, the difference between (A) and (B). Only regions with decoding significantly different from zero are shaded (assessed using two-sided one-sample T-tests corrected using false discovery rate; *q <*= 0.05). Further details in Methods.

### Widespread modulation of cortical activity by uncertainty and change-point probability

Our analyses above suggested that uncertainty and CPP have a marked impact on all readouts used in this study: they are consistent with the idea that both variables (i) drive pupil responses, (ii) modulate the impact of evidence samples on the internal decision, and (iii) also modulate the encoding of the evidence samples in multiple cortical areas. In a final set of analyses, we asked if and where these two ‘modulatory’ variables were decodable from patterns of cortical population activity (Methods). We found this to be the case in an even more widespread manner than for the decoding of evidence and belief states discussed above (Figure 7), suggesting a neuromodulatory signal broadcasting these variables across the entire cerebral cortex to shape the state of local circuits on the fly (see Discussion).

**Figure 7:**
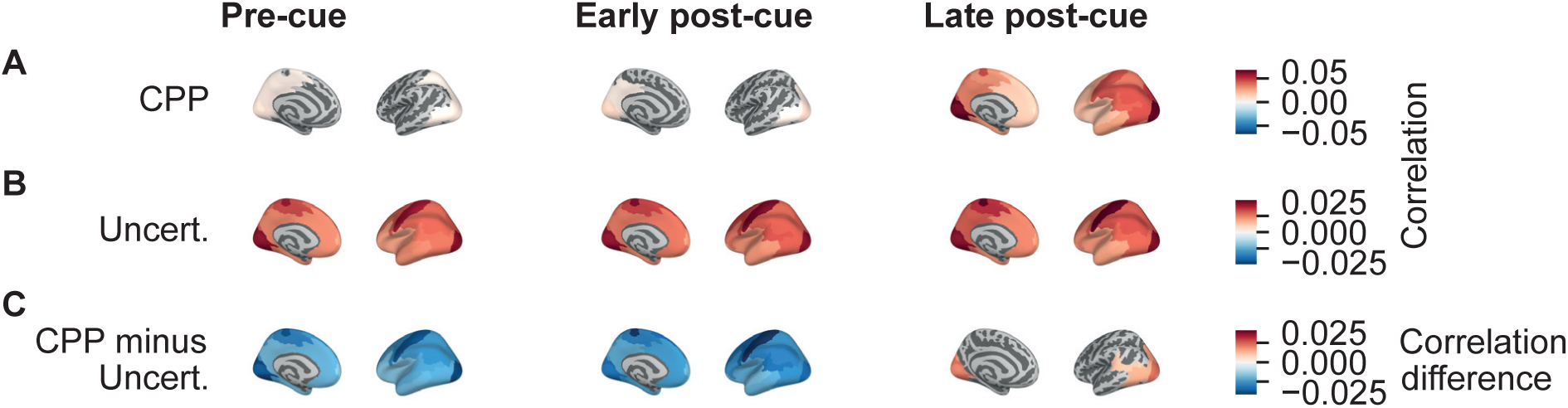
Decoding of change point probability (CPP) and uncertainty relative to the onset of cues in the Inferred condition. Decoding was performed and evaluated analogously to that in Figure 4, using estimates of source activity without time-frequency analysis applied. (A) Decoding of CPP. (B) Decoding of uncertainty. (C) The difference in decoding performance for CPP and uncertainty. That is, the difference between (A) and (B). Only regions with decoding significantly different from zero are shaded (assessed using two-sided one-sample T-tests corrected using false discovery rate; *q <*= 0.05). Further details in Methods.

### No evidence for sensory-motor mapping rule shaping correlated variability of MEG signals

We also tested a pre-registered hypothesis for how the active context would shape intrinsic correlated variability of population codes for the task-relevant inputs and outputs (stimulus orientation and motor action), which was based on a previous fMRI study by van den Brink et al. (2023). We found no evidence for such an effect, neither tested directly (Figure 8, see Methods), nor in an indirect manner (Figure 8; Table 2; Methods). (Although preregistered control tests were significant; Figure 8; Table 2; see Methods). In sum, our preregistered MEG tests were not borne out in the current MEG data.

**Figure 8:**
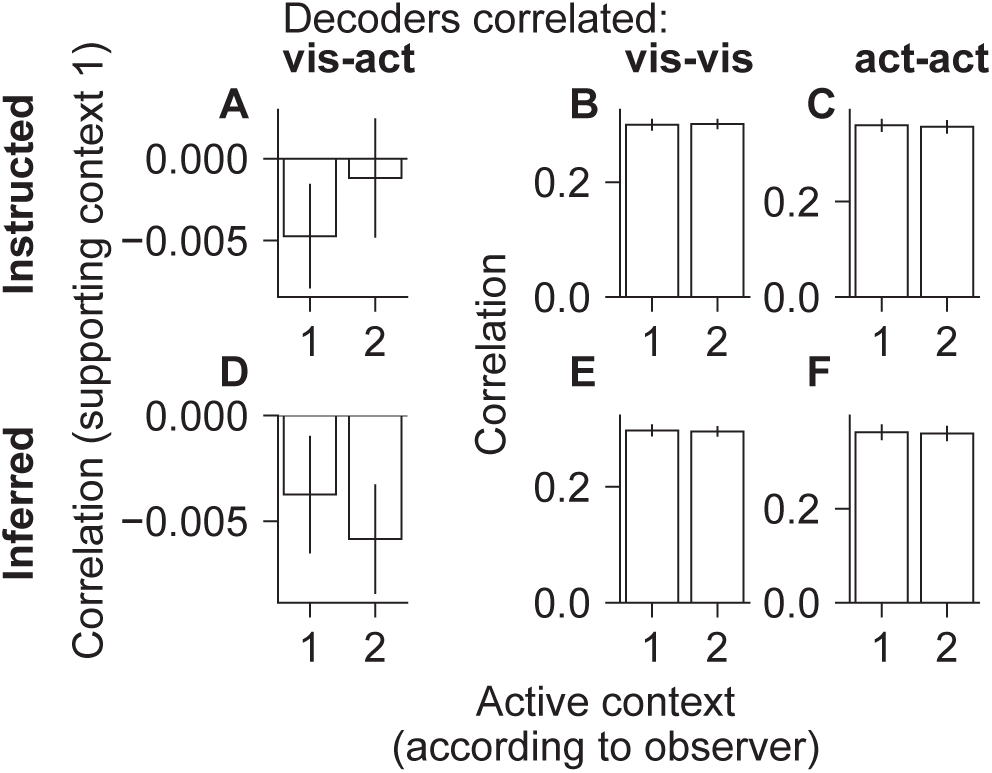
The preregistered comparisons of the correlated variability analysis. Our primary prediction was that when a specific context was active, spontaneous fluctuations in activity in visual representations would be associated with fluctuations in activity in motor representations, in a manner that mirrored the stimulus-to-response mapping rule in that context (see Methods; van den Brink et al., 2023). We measured the latter by training decoders on visual and motor representations and correlating the outputs of visual decoders (“vis”) and action decoders (“act”). (A and B) Positive correlations indicate correlated variability consistent with Context 1, and negative values indicate correlated variability consistent with Context 2. (A) In the Instructed condition we did not observe the predicted difference between Context 1 and Context 2 in decoder output correlations (one-sided permutation T-test, *N* = 19*, T* = *−*1.48*, p* = 0.922), nor were the correlation values positive under Context 1 or negative under Context 2 (Table 2). (D) Similarly, in the Inferred condition we did not observe the predicted difference between the two conditions (one-sided permutation T-test, *N* = 19*, T* = 1.35*, p* = 0.0962), nor positive values under Context 1 or negative values under Context 2 (Table 2). A preregistered control analysis involved looking at the correlation between pairs of visual decoders and pairs of action decoders. In such pairs, both decoders are trained to decode the same target, so we expected the outputs to be positively correlated under all contexts and conditions (see Methods). These preregistered control tests were all significant (one-sided permutation T-tests against zero; *p <*= 0.05; Table 2).

## Discussion

We studied the algorithmic and neural basis of higher-order internal decisions about behavioral context, formed under multiple sources of uncertainty. We demonstrated that human participants apply approximately normative strategies to compute such decisions, and that pupil-linked arousal systems track important latent variables potentially involved in this computation (in particular CPP). Pupil-linked arousal, in turn, may modulate network dynamics across the cerebral cortex, in line with the widespread decoding of CPP from cortical population activity. Most importantly, we uncovered representations of the decision variable (DV) for such internal decisions in many areas of the posterior part of the cerebral cortex, including not only associative regions of posterior parietal cortex, but even early visual cortex. Recent studies mapping the large-scale dynamics of elementary sensory-motor decisions found a similarly widespread distribution of neural population signals encoding accumulated sensory evidence and choice (Khilkevich et al., 2024; Murphy et al., 2021; Wang, 2022; Wilming et al., 2020). In such lower-level decisions, the neural decision variables are commonly encoded in the format of movement plans (Chen et al., 2024; Gold & Shadlen, 2000; Hanks et al., 2015; Khilkevich et al., 2024; Vyas et al., 2020; Wyart et al., 2012). Correspondingly, these decision variables are strongly expressed in populations of neurons that are selective for planning the respective actions: premotor and primary motor cortex for decisions reported by hand movements (Donner et al., 2009; Murphy et al., 2021; Peixoto et al., 2021; Wilming et al., 2020), parietal, frontal and subcortical regions involved in eye movement control for tasks involving saccadic choice reports (Gold & Shadlen, 2007; Kiani et al., 2014), and basal ganglia for several types of action (Ding & Gold, 2013; Khilkevich et al., 2024). In contrast, in our study the decision variable was not associated with any specific movement, but rather informed the internal selection of context for a lower-level decision. This DV was encoded across widespread cortical areas, however less prominently in frontal cortical regions involved in the control of motor action.

Our current results indicate that posterior parietal cortex, including inferior parietal regions that do not contain topographic maps of the visual field (Silver & Kastner, 2009), as well as visual cortex are involved in the computation of a decision variable that governs the internal selection of the stimulus-response mapping used for reporting a low-level sensory-motor decision. While this may be expected for association (i.e., posterior parietal) cortex, the delineation of a robust and stable code for such a high-level decision variable within area V1 is noteworthy. We note that recent animal physiology work has established robust encoding of sensory-motor mapping rules in neuronal response patterns in monkey visual cortex (Jonikaitis et al., 2025) and robust encoding of prior beliefs in neuronal response patterns in mouse visual cortex (Findling et al., 2025). Our current human data are in line with these animal findings and point to a general role of visual cortex in maintaining representations of high-level contextual variables. It is natural to think of the visual-spatial format of the sensory evidence for the higher-level decision as a possible source. Our gaze control analyses refute the possibility that the V1 signals are simply due to retinal shifts of the stimulus display induced by eye movements. A more plausible explanation is that beliefs about the context are propagated to V1 through selective feedback from higher-tier regions. The observation of expression of the belief code in the alpha-frequency band, that has often been associated with cortical feedback is consistent with this idea (Bastos et al., 2015; Mejias et al., 2016; Murphy et al., 2021; Wilming et al., 2020). Whether or not such feedback resembles the mechanism of visual-spatial selective attention (van Kempen et al., 2021; Xia et al., 2024) or uses a different format is an important question. Some evidence supports the notion that the decision signals in early visual cortex can be expressed in non-spatial formats (Nienborg & Cumming, 2009; van den Brink et al., 2023; Wilming et al., 2020).

The presence of high-level decision variables in visual cortex, potentially due to feedback, raises the interesting question of the purpose of these representations. One speculation is that decision-variables are present in sensory cortex because the brain performs approximate Bayesian inference through some form of message-passing algorithm (Bishop, 2006; Findling et al., 2025; Friston, 2010; Friston & Kiebel, 2009; Jardri & Denève, 2013; Minka, 2005; Rao & Ballard, 1999). In such algorithms messages are passed between different nodes, with each node being responsible for performing some part of the inference and sending updated messages to other nodes, necessitating feedforward and feedback connections.

Based on influential accounts (Miller & Cohen, 2001), one might expect robust rule codes in the prefrontal cortex. Our current results do not support this expectation. There are at least two potential reasons for the weak rule decoding in prefrontal cortex. First, rule codes may be robustly expressed in prefrontal cortex, but in a format that is challenging to detect using our source-level MEG approach. For example, neurons encoding opposite rules may be intermingled at a fine spatial scale in this cortical region, diluting any mesoscale biases (of individual vertices or principal components) that our current decoders were using. Indeed, previous fMRI work found only weak evidence for the encoding of (beliefs about) sensory-motor mapping rules in prefrontal cortex (van den Brink et al., 2023; Zhang et al., 2013). Second, rule codes may, at least in the context of our task, be genuinely weak in prefrontal cortex — weaker than in the more posterior regions, where we find strong evidence for such codes. Cellular-resolution data are required to evaluate these possibilities, and are currently lacking for our hierarchical decision-making task.

In a task similar to the current one, van den Brink et al. (2023) found that spontaneous fluctuations of visual stimulus-encoding patterns in multi-voxel fMRI activity in visual cortex, correlated with fluctuations of action-encoding patterns in downstream areas. The structure of this correlated variability flipped with the stimulus-response mapping for reporting a low-level visual orientation judgment (in both the Instructed and Inferred conditions). We predicted that we would find the same context dependence of the structure of correlated variability in stimulus- and action-encoding patterns, also when using source-reconstructed MEG activity. However, we found no evidence for such an effect in the current MEG data. This null-effect could be due to different reasons. One possibility is that fMRI may simply be better suited for tracking spontaneous fluctuations of feature-specific population codes (but see Liu et al., 2022). Another, more likely possibility, is that the current null effect is due to potentially critical differences in the dynamics of rule switches between the tasks used here and by van den Brink et al. (2023): whereas in both conditions of the current study, changes in the active rule were frequent (hazard rate of 8%, Figure 1), the task used by van den Brink et al. (2023) featured much longer stable periods where the rule didn’t change. The hazard rate in their inferred condition was only 1.43%, and in their instructed condition the rule changed in a deterministic and predictable fashion, with long periods of stability due to long delay and ITI periods. Indeed, in that previous study, the strength of the coupling between stimulus and action codes gradually built up over several seconds following the rule switches (see Figure 6 in van den Brink et al., 2023). This is consistent with an involvement of plasticity mechanisms in these stimulus-response associations (Fusi et al., 2007; Miller & Cohen, 2001), which in turn may shape the structure of correlated variability. Our current design may have been too fast for sufficiently strong rule-specific correlation patterns to emerge.

Our pupil results and widespread decoding of uncertainty and change-point probability point to a possible role of diffusely projecting neuromodulatory brainstem systems in broadcasting these latent variables. This idea is supported by the observation in our current study, as well as previous ones (Filipowicz et al., 2020; Murphy et al., 2021; Nassar et al., 2012), that CPP and uncertainty drive pupillinked arousal responses. Brainstem nuclei involved in the control of pupil diameter and central arousal state (Joshi & Gold, 2020; Maheu et al., 2025) may receive information about such high-level computational variables via top-down projections from higher-tier cortical regions involved in uncertainty monitoring (Aston-Jones & Cohen, 2005). The widespread expression of these variables in neural population activity across cortex is also consistent with the widespread modulation of cortical dynamics from subcortical input reflecting change-point probability (more generally, surprise) or uncertainty (Aston-Jones & Cohen, 2005; Dayan & Yu, 2006). However, our current decoding analyses cannot distinguish between cortical representations of these variables (e.g., in areas involved in uncertainty monitoring) and modulations of cortical representations by these variables (e.g., mediated by the resulting neuromodulatory input). Future work could do so by combining our approaches with manipulations of neuromodulator receptors. The locus coeruleus-noradrenaline system has been linked to surprise and uncertainty (Dayan & Yu, 2006; Nassar et al., 2012), but given the relationships between the neuromodulatory nuclei, other systems may be equally involved (Maheu et al., 2025; Sara & Bouret, 2012).

An important question for future research is whether and how our current results generalize to non-spatial and non-visual tasks. A plausible explanation for the occurrence of belief signals in even early visual cortex is the fact that the context evidence in our task is conveyed by visual-spatial stimuli. If we ran the same task but provided the context evidence in a different sensory domain, or in a symbolic fashion (e.g., through numbers), we might expect to find belief representations in the early visual cortex to disappear. At the same time, some of the regions in association cortex identified here, specifically those in inferior parietal cortex, which lack clear visual field maps (Silver & Kastner, 2009), encode higher-level decision variables for internal decisions in an abstract format that does not depend on the input channel conveying the evidence.

The behavioral model we considered does not take into account computational costs (Tavoni et al., 2022). In real life, humans and other animals are not likely to track all variables all the time as prescribed by the normative solution, because this is effortful (i.e., costly). CPP and uncertainty, the two latent variables which we found to drive pupil-linked arousal and dynamically modulate the weighting of evidence for the internal context decision, may be more important for resource-optimal cognition than postulated by the current cost-free normative model: These variables may not just up-weight evidence in a graded manner, but dynamically control when to commit limited computational resources (O’Reilly et al., 2013; Tavoni et al., 2022). A related consideration is that our current results do not allow us to pinpoint specific roles of the brainstem arousal systems driving the pupil responses to tasks such as ours (Joshi & Gold, 2020): arousal may modulate the level of internal noise (Aston-Jones & Cohen, 2005; Pfeffer et al., 2021), up-weight the impact of the new evidence relative to the prior (Jordan & Keller, 2023; Murphy et al., 2021; Nassar et al., 2012; Yu & Dayan, 2005), or control network reset to allow new stimulus-response mappings – or, more generally, new strategies – to be formed (Bouret & Sara, 2005; Fusi et al., 2007). Disentangling these different alternatives will be an important goal for future work.

Our results suggest that humans accumulate evidence in an adaptive fashion, even when the resulting decisions are not directly associated with a motor action. To achieve this, key quantities such as surprise (more precisely, change-point probability) or uncertainty are computed, tracked, and may be broadcast by neuromodulatory systems of the brainstem. The evolving belief about an internal decision is itself widely represented across the posterior cortex, including even early visual cortex.

## Methods

### Participants

19 healthy humans (10 Female, 9 Male, 0 Non-binary, 0 Prefer not to say) participated in the study, which was approved by the ethics committee of the Hamburg Medical Association (Ärztekammer Hamburg). Participants were recruited through posters and an established mailing list. A target of 24 participants was initially set because this would lead to a well-balanced randomization of stress/pharmacological conditions described below, which are not analyzed in the current study. This target was reduced to 20 participants, due to a large backlog of planned MEG studies after a long Covid-induced measurement pause and, consequently, limited MEG time allotted to each individual study. Due to several cancellations, we ultimately arrived at a sample size of N=19, which is well within the range of previous successful MEG studies (e.g., Murphy et al., 2021; Wilming et al., 2020). A power analysis suggested that the obtained sample size was sufficient to test the preregistered pupil hypotheses (see pupil-analyses preregistration; https://osf.io/hnwfr). Importantly, we collected a large amount of data from each participant after extensive practice in the task (1632 non-training trials per participant), which had a higher priority for in-depth modeling of individual behavior and in-depth individual MEG data analyses.

Participants were paid approximately 411 EUR for approximately 14 hours participation distributed over 7 sessions, including: informed consent and behavioral practice, main MEG experiment (4 sessions), informed consent and clearance for structural MRI, and structural MRI scan. An extensive set of inclusion and exclusion criteria were applied. This included the following: participants were between 18 and 40 years old, had no physical, neurological, or psychiatric illnesses, no relevant allergies, were not pregnant, and had not taken medication in the previous two months. Participants were also excluded if they had MRI incompatible objects in their body, or if they had insufficient visual acuity (without glasses) at a 60 cm distance.

### Experimental design and general procedure

In a first session, each participant received information about the study and task instructions, and then practiced the task in a behavioral psychophysics laboratory. Each participant underwent four main experimental sessions, all at the same time of day, within the MEG laboratory. At the beginning of each main experimental session either 0.5 mg prazosin (*α*1-receptor antagonist) or a placebo (visually identical to the drug) was administered orally, in a randomized, and double-blind design. Each participant took prazosin twice and the placebo twice. The first non-training trial of the main behavioral task started at least 55 minutes after drug (or placebo) administration. In addition, a stress manipulation, the “cold pressor test” (hand immersion in ice water), or a corresponding control condition (hand immersion in moderately warm water), was applied (Hines & Brown, 1936; Schwabe et al., 2008). These conditions (“ice water” vs. “warm water”) were blocked by session, and repeatedly applied within each session, initially prior to the first main experiment block, and then repeated every two blocks, giving a total of 6 immersions per session. Participants were asked to keep their hand in the water for as long as they could, up to 3 minutes. Within each of the 4 main experimental sessions, measurements of blood pressure, heart rate, and self-reported stress were performed. Pharmacological and stress conditions were coupled: Administration of prazosin was always linked to the active stress condition (“ice water”) and placebo was always linked to the control stress condition (“warm water”). The coupling of the pharmacological and stress conditions was the result of a technical error, and was for this reason not specified in the preregistration. Importantly, behavior was qualitatively and quantitatively similar in the “ice water” and “warm water” conditions: Figure 2 – Supplemental Figure 3 shows the results of the regression performed in Figure 2 but now including separate effects for the “ice water” condition (see below). There were no significant differences between the conditions for the effect of LLR, or the modulatory effect of CPP. Only at a single lag for the modulatory effect of uncertainty was there a significant difference between the two series. Therefore, all main analyses reported in this paper collapsed across those conditions.

### Behavioral task

#### Apparatus

The behavioral task was presented using MATLAB and Psychtoolbox-3 running on Linux machines at 60Hz (120Hz during training session; Brainard, 1997, Kleiner et al., 2007, Pelli, 1997). During the training session stimuli were presented on a VIEWPixx monitor (approximately 60cm from participants and with a physical size of 52.2 × 29.3 cm), while during the MEG sessions stimuli were presented via projection onto the back of a transparent screen (PROPixx VPX-PRO-5050B projector), which was approximately 60 cm from participants (70 cm for the first four participants) producing an image of size 45 × 26.5cm (43.5 × 25.5cm for the first four participants).

#### Task structure

The task required participants to judge the orientation (horizontal or vertical) of full-contrast visual grating stimuli (presented for 0.6s) and report their judgment by pressing a button using either the left hand or right hand, with the correct stimulus-response mapping defined by the context (see below; Figure 1; van den Brink et al., 2023). The grating stimulus had a radius equal to 45% of total screen height; in the MEG lab its diameter corresponded to 22.5 DVA for most participants, and 18.6 DVA for the first four participants. Button press responses were delivered on two keyboard keys during the training session (behavioral psychophysics laboratory), and two buttons on two different response boxes during main experimental sessions (MEG laboratory). There was no deadline for reporting the judgment. Further, only responses occurring at least 0.1s after the grating stimulus cleared from the screen were counted. The fixation cross rotated at this time point, to indicate to participants that they could now respond. (Early responses led to a warning message that lasted for 1.5s.) The gratings had a central aperture so as not to obscure the fixation cross that was presented throughout each block of trials, including during the inter-trial-intervals (ITIs; radius of aperture was 12% of total screen height). We define a “trial” of the (lower-level) orientation judgment task as the interval ranging from the presentation of a grating stimulus to the subsequent behavioral response of the participant (Figure 1). At the beginning and end of each ITI, and end of each trial, there were also short periods during which only the fixation cross was presented (0 - 0.9s, uniformly distributed, at the beginning of each ITI, and 0.3s at the end of each trial).

The orientation judgments in the lower-level task had to be reported using one of two stimulus-response mappings. Which stimulus-response mapping was required was determined by the current context, which alternated throughout the course of each experimental session and block of trials. Under “Context 1”, participants were asked to report the horizontal orientation with a left-hand button press, and vertical with the right-hand button press (Figure 1). Under “Context 2” the required stimulus-response mappings were reversed. Trial-by-trial feedback was not provided, but at the end of each block participants were informed of the percentage of trials in which they had answered correctly.

At the beginning of each block (of 34 trials) the active context was randomly drawn. Prior to the presentation of each cue there was an 8% probability that the current context changed to the alternative context (i.e., a “hazard rate”, *H* = 0.08; Fig 1). The active context did not change during a trial, only during ITIs.

Information about the current context was provided by a sequence of visual cues (each of 0.15s duration with 0.4s onset asynchrony) presented in rapid succession during the inter-trial-intervals (multiple of 0.4s from 1.2 - 4s inclusive, uniformly distributed). The cues were the locations (more precisely, the polar angles) of small dots presented simultaneously in the left and right visual hemifields at fixed eccentricity (2.02 DVA for most participants and 1.67 DVA for the first four participants). These dots were always presented in pairs in the left and right visual hemifields, at symmetric polar angles mirrored about the vertical meridian. This was done for symmetry of the display, and to minimize the tendency to track individual dots. Dot locations varied across the upper and lower portions of the visual hemifield. Because both dots of a pair provided redundant information (due to the above mirror symmetry), we refer to each pair of dots as one cue. We denote the polar angle (in radians) of the right-hemifield dot, measured in the plane of the display, clockwise from horizontal, for the *n*th cue in the block, by *x_n_*.

The reliability of the information provided by cues about the current context varied between two conditions that we refer to as “Instructed” and “Inferred” (van den Brink et al., 2023). Within each block, one of these two conditions applied. Participants were informed of the upcoming condition prior to the start of each block.

In the Instructed condition, cues were fully informative of the context. If the context was “Context 1”, then the pair of dots flashed at a fixed angle (in the plane of the display) in the lower visual hemifield (12 degrees below horizontal), and if the current context was “Context 2” then the pair of dots flashed at a fixed angle in the upper visual hemifield (12 degrees above horizontal). Because of the fixed, noiseless cue positions, there was no uncertainty about the context in this condition: Participants could determine the context simply from the last cue presented, and select the stimulus-response mapping accordingly.

The Inferred condition functioned in a similar way, but in this condition noise was added to the cue locations, so that the information they provided was no longer completely reliable. Here, the context determined the (truncated) Gaussian distribution from which cues were drawn: if the active context at the *n*th cue, *C_n_*, was *j* (where *j* = 1 indicates “Context 1” and *j* = 2 indicates “Context 2”) then (the untruncated distribution was given by),

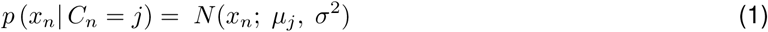

where *N* (*a*; *b, c*) indicates a normal distribution with mean *b* and variance *c*. This distribution was then truncated at the vertical meridian. The standard deviation was fixed (20 degrees), but the mean differed for the two contexts and matched the positions used in the Instructed condition (i.e., ±12 degrees). More explicitly,

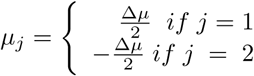

where Δ*µ* parameterizes the difference between the two means. Participants needed to infer the underlying mean in order to select the correct context in any given trial. In other words, any single cue could be misleading, and the optimal (maximizing task accuracy) solution to infer the current context (generative mean) entailed accumulating the information provided by more than just the immediately preceding cue, while taking into account the possibility that the context could, prior to any cue, change.

#### Behavioral task procedure

At the start of each MEG session participants completed 2 training blocks (one Instructed block, followed by one Inferred block), which included trial-by-trial feedback (in main experiment blocks trial-by-trial feedback was not provided). Each session contained 12 blocks of the main behavioral task (ordered using the pattern Inferred-Instructed-Inferred repeated 4 times). The behavioral practice session prior to the MEG sessions featured 8 training blocks, and 6 main behavioral task blocks.

Following the training blocks, but prior to the main experiment blocks, and again after all main experiment blocks were complete, participants completed a “localizer” block (hence there were 2 localizer blocks in total). During these blocks participants were asked to perform a change detection task on the central fixation cross: When the cross began rotating, they were to report the direction of rotation with a button press (left for anticlockwise, right for clockwise). While participants were told that they could ignore everything else presented on the screen, horizontally and vertically oriented gratings with a central aperture – of the same kind used in the main experiment – were periodically presented.

### Data acquisition

MEG recordings (at 1200 Hz) were conducted in a room with magnetic shielding, using a CTF whole-head MEG system with 275 axial gradiometers. During MEG sessions eye tracking data was acquired using an SR Research EyeLink 1000 device (at 1000 Hz). An additional (uncalibrated) version of the eye tracking data was collected with the MEG recordings at 1200 Hz through the use of an SR Research analog card.

The main version of the eye tracking data (recorded at 1000Hz; EyeLink) was calibrated and validated with a 9-point fixation routine. At the start of each session, a reference recording was also conducted using a surrogate pupil of known size.

Throughout MEG sessions head position was recorded using three fiducial coils (two placed using ear plugs in the ears, and one adhered to the skin at the nasion). At the beginning of each session, once the participant was comfortable, the position of the head was recorded. Following this, prior to every block (if required) the participant was given feedback on how to adjust, such that their head was as close as possible to the original position (Stolk et al., 2013).

An electrocardiogram (ECG) was recorded throughout using two electrodes (placed just below the collarbone on the participant’s right side, and just below the ribs on the participant’s left side). Additionally horizontal and vertical electrooculography (EOG) recordings were taken using a pair of electrodes above and below the participant’s left eye, and a pair of electrodes to the left of the left eye and right of the right eye. Finally, a ground electrode was applied above the participant’s left wrist.

Structural MRI measurements for MEG source reconstruction were performed on a 3T Siemens Trio scanner, using a T1-weighted MPRAGE sequence (TR 2, 300 ms; TI 1, 100 ms; TE 2.98 ms; flip angle 9*^◦^*; FoV 256 mm; slice thickness 1 mm; voxel resolution 1 × 1 × 1 mm^3^).

### Preregistrations

We preregistered the following exclusion condition: We would exclude participants from analyses of the respective conditions if they did not achieve a minimum average of 55% accuracy in the Inferred condition, or if they did not achieve an average of 60% accuracy in the Instructed condition. No participants were excluded on this basis. Additionally, no participants were excluded for completing one or more MEG sessions, but not all four.

A subset of analyses were preregistered through two separate preregistration documents, one covering the normative model and pupil analyses (see below; https://osf.io/hnwfr), and the other covering the MEG correlated variability analyses (https://osf.io/h4×5s). Where a preregistration applies this is explicitly noted.

### Behavioral data analysis and modeling

Data analysis was conducted using code written in Matlab (MathWorks) and Python (see “Code availability”). Only data from the main experimental blocks of the MEG sessions were used for the behavioral modeling. We computed overall percentage correct choices (i.e., orientation judgments using the appropriate stimulus-response mapping) for both Instructed and Inferred conditions. When comparing performance in the two conditions we used a paired t-test. We used the one-sample variant of Cohen’s *d* for effect size (Cohen, 1988). This involved computing the difference in performance between the two conditions for each participant, taking the mean of these values, and dividing by their estimated population standard deviation.

All behavioral modeling was performed on the data from the Inferred condition, unless explicitly noted.

#### Normative model

We used a normative model for evidence accumulation in changing environments introduced by Glaze et al. (2015). This model prescribes the Bayes-optimal strategy for making two-alternative forced choice decisions about the active state of a changing environment based on noisy cues and under a fixed level of volatility (hazard rate). Here, we apply this model to the higher-order, internal decision of selecting the context that, in turn, determines the stimulus-response mapping for reporting the lower-order visual orientation judgment.

We assume that participants have understood and learned the general statistical structure of the task when performing the main experiment. This assumption is plausible because (i) the task structure is explained to participants prior to task performance in a simplified manner (e.g. exact probability distributions not described), and (ii) participants had practiced the task extensively in a separate session before the main experiment. Even so, we allow the possibility that participants misestimate certain statistics, specifically the hazard rate and/or the mapping from dot locations to decision evidence. The normative strategy for inferring the state (i.e., selecting the context) is to compute the log-posterior ratio after cue *n*, denoted *L_n_*, by combining the evidence provided by that cue with the prior (before observing the cue) using Bayes rule. The evidence is expressed as a log-likelihood ratio for cue *n*, *LLR_n_*, and the prior is a non-linear transformation of the log-posterior ratio after the previous cue. The required computation is (Glaze et al., 2015):

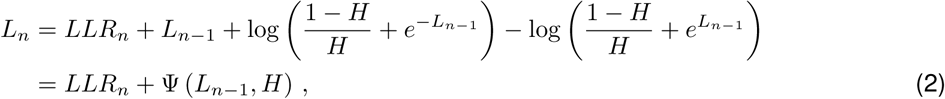

where abbreviation Ψ(*a, b*) stands for the log-prior ratio (Glaze et al., 2015). To simplify equations, we will sometimes use the alternative notation Ψ*_n_* where Ψ*_n_* = Ψ (*L_n−_*_1_*, H*).

The log-likelihood ratio for a cue can be computed using the fact that the *x_n_*are normally distributed (ignoring minor truncation effects), by substituting in the specific normal distribution for the *x_n_* and rearranging. This gives (Glaze et al., 2015; Gold & Shadlen, 2001),

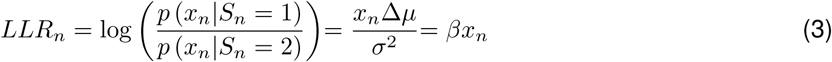

where *β* is an abbreviation. We see that cue values, *x_n_*, (i.e., angles in radians of the right-hemifield dot from horizontal in the plane of the display) are simply scaled by *β* to convert them to log-likelihood ratios, and we therefore refer to *β* as the evidence scaling.

At the onset of each trial of the orientation discrimination task, participants need to select a context they are going to use for their behavioral report. That is, they are forced to perform a two-alternative selection of the active context. The ideal observer would pick the context based on the sign of the log-posterior ratio, *L_n_*, in a deterministic fashion. We allow for the possibility that the participants’ context selection decisions based on *L_n_* are subject to noise. Denote the case in which the observer chooses context *C* = 1 and uses the corresponding stimulus-response mapping by *R* = 1. Similarly, denote the case in which they choose *C* = 2 by *R* = 2. In the model these decisions follow,

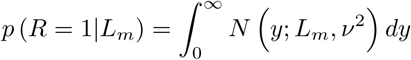

where *m* is the number of the final cue before the trial onset, and *ν* parameterizes the standard deviation of the decision noise. The probability for the alternative decision can be calculated by noting that the probability of both decisions must sum to 1.

We use three variants of the normative model, which differed in their assumptions about how well participants represented the statistics of the task. In the basic normative model (“Normative”) we assume that participants correctly estimate both the hazard rate, *H*, and the evidence scaling, *β*. The standard deviation of the decision noise *ν* is the only free parameter. In the “Miscalibrated h normative” model we allow the possibility that the observer misestimates the true hazard rate and instead uses *Ĥ* in Eq. (2), where *Ĥ* is fit as a free parameter (in addition to *ν*; two free parameters). Finally, in the “Miscalibrated h and b normative” we additionally allow the possibility that the observer does not use the correct evidence scaling, *β*, but some other value *β̂*, for converting dot locations into decision evidence (*LLR*, Eq. (3)), yielding a total of three free parameters.

#### Alternative models

We also considered simpler heuristics that differed qualitatively from the normative strategy (Figure 2 – Supplemental Figure 1; Glaze et al., 2015). In the “Last sample” model, the observer only uses the final cue prior to the forced-choice to make their decision (i.e. the final evidence sample). Specifically, they use the angle from horizontal of the cue in radians. Analogously to the normative model, decisions are based on a noisy version of this decision variable. This gives one free parameter, the standard deviation of the normally distributed decision noise, *ν*.

In the “Perfect accumulator” model the observer “perfectly” accumulates all evidence presented, by keeping track of the sum of the dot locations (in radians) of all cues presented since the start of the block (Bogacz et al., 2006). Again, decisions are based on a noisy version of this decision variable (one free parameter: decision noise).

The “Bounded accumulator” model is similar to the “Perfect accumulator” model, except that the accumulator integrating the evidence samples cannot exceed a value Θ, or go below *−*Θ, which acts as a non-absorbing bound (Glaze et al., 2015). For example, if the accumulator is at Θ and the observer sees a cue providing positive evidence, the accumulator simply remains at Θ, instead of increasing beyond this value. The observer is again subject to decision noise, meaning this model has two free parameters: *ν* and Θ.

Finally, the “Leaky accumulator” is another variant of the “Perfect accumulator” in which the observer again integrates the evidence samples but now, prior to each update to the accumulator, some fraction of the previously accumulated evidence is discarded (Glaze et al., 2015; Ossmy et al., 2013; Usher & McClelland, 2001). This is achieved by multiplying the state of the accumulator by (1 *− α*). If *α* = 1, all previous information is lost, making the model identical to the “Last sample” model. If *α* = 0, no previous information is lost, making the model identical to the “Perfect accumulator”. The “Leaky accumulator” has two free parameters, *α*, and again a parameter for decision noise, *ν*.

#### Model fitting

We fit the models on a participant-by-participant basis, by maximizing the log-likelihood of participant’s responses using MATLAB’s “fmincon” optimization algorithm (to be precise the negative log-likelihood was minimized; ‘Matlab Optimization Toolbox’, 2017). The upper and lower bounds for the parameter values during fitting are shown in Table 1.

**Table 1:**
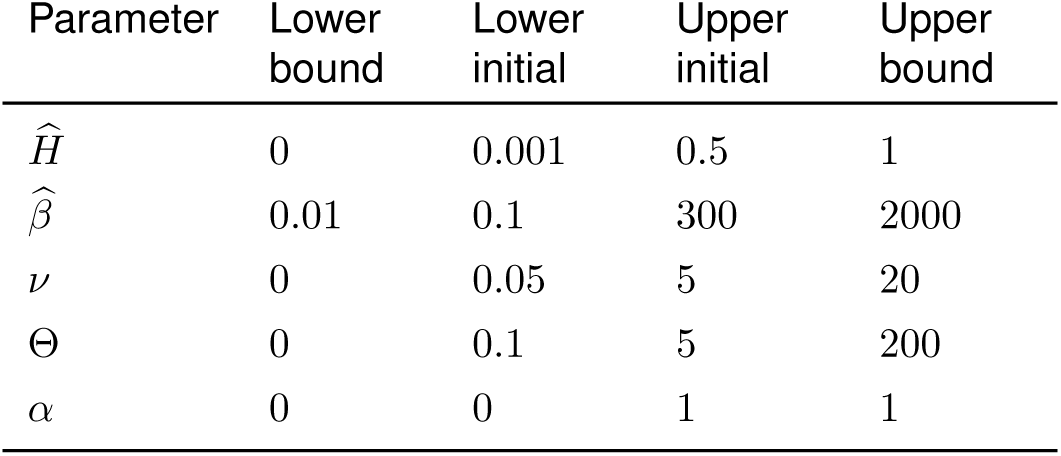
Bounds placed on model parameters during fitting, along with the initial range from which candidate start points for the fitting were drawn.

| Parameter | Lower bound | Lower initial | Upper initial | Upper bound |
| --- | --- | --- | --- | --- |
| $\hat{H}$ | 0 | 0.001 | 0.5 | 1 |
| $\hat{\beta}$ | 0.01 | 0.1 | 300 | 2000 |
| $\nu$ | 0 | 0.05 | 5 | 20 |
| $\Theta$ | 0 | 0.1 | 5 | 200 |
| $\alpha$ | 0 | 0 | 1 | 1 |

We began each fit by drawing 250 sets of candidate starting points for the optimizer, where a candidate “starting point” contains a candidate value for each parameter. Candidate values for parameters were drawn from uniform distributions on the interval between the “Lower initial” and “Upper initial” shown in Table 1. The candidate starting point with the greatest log-likelihood was then used as the optimization start point.

This model fitting procedure was rerun 10 times for each participant, and the maximum log-likelihood found over all fits, and corresponding parameter values, were used for all further analyses. Repeated fits minimize the risk of only discovering a local maximum of the log-likelihood function and allow for some assessment of associated issues (Acerbi et al., 2018, supplementary methods).

#### Model-based latent variable estimation

We then extracted the log-posterior ratio and log-likelihood ratio after each cue using Eq. (2) and Eq. (3) from the fitted “Miscalibrated h and b normative” model, which were used for several of the MEG analyses described below. In some analyses, we also use additional latent variables derived from the normative model. Specifically, we computed a simple measure of the observer’s uncertainty prior to observing cue *n* using *−*1 times the absolute value of the log-prior ratio (Murphy et al., 2021):

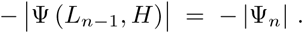

We also calculated “change point probability” or CPP, the posterior probability that the state changed between cue *n* and cue *n −* 1, given the previous belief state *L_n−_*_1_ and the new sample of evidence, through (Brink et al., 2024; Murphy et al., 2021):

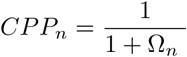

where,

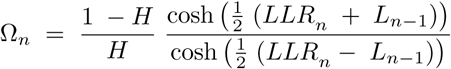

For most analyses we computed these variables on a cue-by-cue basis using the fitted parameter values for *H* and *β* from the “Miscalibrated h and b normative” model (i.e. replacing *H* and *β* with *Ĥ* and *β̂* in all computations of the latent variables). Where explicitly stated we used the true values instead.

#### Model comparison and data simulation

We compared the goodness of fit of the various models using Akaike information criterion (AIC) and Bayesian information criterion (BIC) values calculated for each participant and model. Separately for each information criterion, we found the overall best fitting model. We then computed, on a participant-by-participant basis, relative information criterion values, by subtracting off the value of the information criterion for the best fitting model. We compared these relative information criterion values to zero, in order to test whether the difference between each model and the best fitting model was significant. For this purpose we computed 95% confidence intervals via bootstrapping (10000 draws).

Additionally, we examined how well the fitted models captured diagnostic qualitative features of the Inferred condition behavioral data. To achieve this we simulated data from the fitted models. Our standard approach was to simulate new stimuli (in the same way the real stimuli were generated), and then to simulate responses based on the model and fitted parameters in question. We simulated the same number of participants as the real number of participants (and used the fitted parameters for each real participant to generate the data for the corresponding simulated participant). In order to reduce noise in the simulations we simulated more trials than in the real experiment (340 trials per block).

To save computation time, when simulating data from all the considered models for the comparison of accuracy across models in Figure 2 – Supplemental Figure 1, we used the stimuli that were actually presented to participants (and hence the number of simulated trials matched the number of trials used in the experiment).

The “theoretical maximum” performance used in Figure 2 was also estimated via simulation of optimal calibrated observers, not subject to any form of internal noise. We simulated a dataset with the same shape as the real dataset, except we simulated 2000 trials per block.

#### Exploring features of the data

To explore the effect of LLR, LLR × CPP and LLR × uncertainty in Figure 2 we conducted a similar analysis to the regression-based analysis performed by Murphy et al. (2021). We predicted the stimulus-response mapping (i.e. context) used by participants in the Inferred condition in a logistic regression. The variables used were LLR, CPP, and uncertainty (*− |*Ψ*_n_|*) associated with the cues leading up to the response (we considered the 15 previous cues), along with an intercept term:

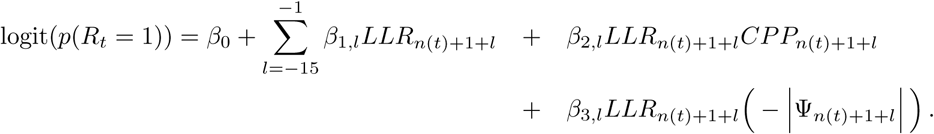

Here *t* gives trial number, and *n*(*t*) is the index of the cue immediately preceding trial *t* (i.e., *LLR_n_*_(_*_t_*_)_ is the log-likelihood of the final cue presented before the response under consideration). *p*(*R_t_* = 1) is the probability that the participant makes a response consistent with Context 1. *l* gives cue number relative to the response (with *−*1 indicating the immediately preceding cue). The various *β* coefficients are the regression weights (and are unrelated to the evidence scaling parameter of the normative model). All predictors were individually z-scored prior to conducting the logistic regression (not explicit in the above equation). We used the true (not fitted) parameter values in the normative model to compute LLR, CPP, and uncertainty. Following the regression, we binned the regression weights on the basis of the cue number relative to response that they corresponded to (i.e. on the basis of *l*) separately for the LLR weights (*β*_1_*_,i_*), the CPP modulation weights (*β*_2_*_,i_*), and the uncertainty modulation weights (*β*_3_*_,i_*). Bins contained 3 weights, and we averaged the weights within each bin.

We also performed a variant of this analysis that included effects of the cold pressor test (Figure 2 – Supplemental Figure 3). This analysis was conducted in the same way as before, except that we now generated a dummy variable (1 or 0) for the use of ice water (i.e. use of the cold pressor test), *I_s_*_(_*_t_*_)_. Here *s*(*t*) gives the session to which trial, *t*, belonged. We then included additional terms reflecting the interaction between this dummy variable and the previously considered variables. That is, the following terms were included in the summation over *l*:

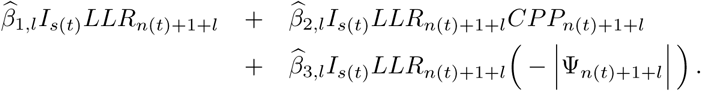

The various *β̂* coefficients are the additional regression weights (and are again unrelated to the scaling parameter of the normative model). This allowed us to plot previously considered effects of LLR, CPP and uncertainty, but now both on cold pressor test sessions (“ice water”), and sessions without the cold pressor test (“warm water”).

Log-likelihood values used in the regression analysis for Figure 2 are correlated with neighboring values, due to the fact that cues close to each other tend to be from the same context, and therefore provide similar evidence. We also performed an evidence “residuals” analysis on Inferred condition data (Figure 2 – Supplemental Figure 2) to deal with this (Charles & Yeung, 2019; Resulaj et al., 2009). In this analysis we computed evidence residuals by, for each cue, subtracting off the mean value of the distribution from which that cue was drawn (i.e. the mean of the distribution in Eq. 1). This leaves us with evidence residuals that are no longer correlated with each other (because the residuals correspond to independent noise added to the mean cue positions). For each response, we then consider the immediately preceding cues, and multiply these cues by 1 or *−*1 so that they no longer give evidence in favor of Context 1, but rather evidence in favor of the context used by the participant in that trial. We then have a set of values associated with each response, and averaging across responses we are left with the average evidence residuals favoring the context used. Evidence residuals at time points that were used to inform the decision will, on average, be consistent with the decision made. Hence, average evidence residuals in the direction of the context used will be positive at such time points. Evidence residuals at time points that were not used to inform the response are random values, randomly signed, and so will average to zero (Resulaj et al., 2009).

Both the evidence residuals analysis (Figure 2 – Supplemental Figure 2) and regression analysis (Figure 2) were performed on a participant-by-participant basis. The corresponding plots show average values across participants, along with error bars of *±*1 standard error of the mean (SEM) across participants. Significant points (across participants at *p <*= 0.05) were computed using permutation cluster-based t-tests with threshold-free cluster enhancement (as implemented by MNE; neighboring points in time considered adjacent; Gramfort et al., 2013; Larson et al., 2024; Maris and Oostenveld, 2007; Smith and Nichols, 2009).

### Pupil data analysis

#### Preprocessing

Pupil analyses focused on the Inferred condition data. We used the main version of eye tracking data (collected at 1000Hz on the PC running the EyeLink eye tracker). Individual pupil recordings for which > 50% of data points were identified as artifactual (see below) were excluded from the pupil analysis.

In the pupil diameter time series blinks and missing data segments were linearly interpolated across (de Gee et al., 2017; Knapen et al., 2016). We identified other artifacts by the derivative of the pupil diameter exceeding a threshold of 25 pixels and interpolated across these (van den Brink et al., 2016, 2023). This process was repeated 100 times to ensure all artifacts were identified. All sections that were thus identified to contain artifacts were then also interpolated across in the gaze x and y position data.

Following interpolation, additional artifacts in a 1 s period preceding the onset and following the offset of blinks / missing data were first estimated using deconvolution and then removed via multiple linear regression (Knapen et al., 2016).

The associated preregistration referred to conversion to mm, but this procedure has no effect, given the following z-scoring (which was also preregistered). The pupil time series was z-scored prior to regression, yielding a time series of z-scores referred to as *Pupil*(*τ*) in the following. (If using the derivative of pupil diameter, z-scoring was conducted after the derivative was taken.)

#### Regression analysis

Following Murphy et al. (2021), the relationship between the pupil and behavior was examined using linear regression with latent computational variables as predictors. Latent computational variables were computed on a participant-by-participant basis using the variant of the normative model in which hazard rate (*H*), evidence scaling (*β*), and decision noise (*ν*), had all been fit. The outcome of the regressions were either pupil diameter, or its derivative. The regression (conducted separately for each participant) had the following form:

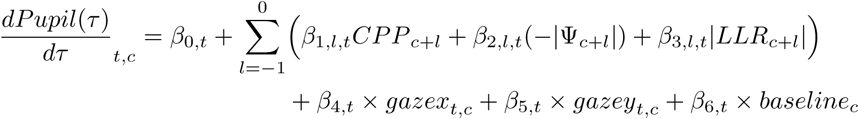

Where *t* is time relative to cue onset, *c* indexes cues, 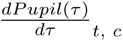 gives the value of the pupil derivative time series at time *t* relative to the onset of cue *c*. CPP is change point probability, *−|*Ψ*_n_|* is uncertainty, LLR is the sensory evidence, gazex and gazey are the x and y gaze positions on screen (indexed in the same manner as the pupil derivative time series), and baseline is average pupil diameter in a 100 ms window surrounding cue onset. The terms relating to prior cues are included to account for auto-correlation in the pupil response and to isolate responses to the current cue. However, these terms were left out in the event that they resulted in a collinearity problem in the regression model (variance inflation factor > 10). Due to a mistake in the preregistration, the regression described here differs slightly from the regression described in the preregistration: Here all beta terms have a *t* subscript, instead of just some of them. We only ran the analysis as described here.

The results for this base analysis are shown in Figure 3 – Supplementary Figure 1. In the main text we present the results for two additional variants of this regression model. First, in order to exclude that any relationship between pupil and CPP was driven by low-level visual differences between consecutive cues, we also ran a variant of this regression model that included one additional term representing “sensory difference”: *|LLR_c_ − LLR_c−_*_1_*|* (“CPP” analysis in Figure 3; Wyart et al., 2012). Second, we fit a version of this regression model to baseline corrected pupil diameter (rather than the derivative; “Uncertainty” analysis in Figure 3). This later regression model did not include baseline diameter, or terms for preceding cues.

Based on prior work (Murphy et al., 2021), we expected CPP following cue onset to co-vary with the derivative of pupil diameter, and we expected uncertainty to co-vary with pupil diameter. To test these hypotheses we compared the resulting regression coefficients to chance (zero) using non-parametric permutation testing (within-participant shuffling with 10,000 iterations; shuffling was conducted across time points simultaneously). Significance was assessed after correction for multiple comparisons across time points using the false discovery rate (FDR; *q <*= 0.05). These tests were preregistered.

For the effects of uncertainty on pupil diameter (Figure 3), no points were significant using the preregistered test. We ran an additional, more powerful statistical test (not preregistered), that took into account the temporal structure of the data. Namely, we used the same permutation cluster-based t-tests used for the regression analysis exploring the effects of LLR, LLR × CPP and LLR × uncertainty (described above; neighboring points in time considered adjacent).

#### Average y-gaze

For the histogram of gaze position in Figure 4 – Supplemental Figure 3 we computed average positions of the gaze along the y-axis of the display (i.e. average position along the vertical axis). Data were preprocessed as before but without the final z-scoring step.

Separately for each participant and separately for cues in which the prior (immediately before seeing the cue) supported context 1 vs. context 2, we computed average y-gaze position in degrees of visual angle (DVA) in the “Pre-cue” time window that we also use in the decoding analyses below (*−*0.05 *<*= *t <*= 0.025 s relative to cue onset). This resulted in two average values for each participant, which were then plotted in the corresponding histogram in Figure 4 – Supplemental Figure 3.

#### Preregistration

As described above, a subset of tests were preregistered. The pupil analysis preregistration was prepared and deposited with the OSF during data collection (https://osf.io/hnwfr). At the time of preregistration some aspects of the data had been accessed for ensuring all required data was being collected and was of satisfactory quality (e.g. MEG, behavioral and eye tracking data had been visually examined, and plotting of response time and cue position distributions had been conducted). Due to the timeline of linked projects it became necessary to conduct some limited analysis prior to preregistration: The behavioral model had been fit to the data for the purpose of estimating latent variables (described above). At the time of preregistration no analyses using these latent variable estimates had yet been conducted. Code for preprocessing of pupil data had been written and tested on the first block of the first participant, but not the remaining blocks or participants.

### General MEG data analysis

MEG analyses used a mixture of tools from FreeSurfer (Fischl, 2012), MNE toolbox code (Gramfort et al., 2013, 2014; Larson et al., 2024; https://mne.tools/stable/index.html), and code from previous projects from our laboratory (Murphy et al., 2021; Wilming et al., 2020; https://github.com/DonnerLab/pymeg; https://github.com/DonnerLab/meg-preproc) in Python.

#### Preprocessing

Preprocessing of MEG data was performed on a session-by-session basis. Visual inspection revealed a number of channels to be clearly defect. These channels were removed (this step differs from the preregistered analysis). An automated algorithm was used to exclude segments of the MEG data where the signal from a sensor was flat for more than 5 ms. Sensors where more than 5% of the data was flat were excluded, rather than excluding the corresponding timepoints for all sensors. Line noise was removed with a notch filter (zero-phase finite impulse response; FIR; zero-phase achieved by using a linear-phase filter and compensating for the delay; Gramfort et al., 2013, 2014) designed to filter out 50 Hz and multiples of 50 Hz up to and including 400 Hz.

Only for the purposes of conducting an independent components analysis (ICA), the data were then high-pass filtered at 1 Hz (zero-phase FIR). Following this, only data from main experiment blocks and localizer blocks were retained. Additional sections of data were excluded (again for the ICA only) by dividing the data in 1 second segments. Any segment was excluded for which, on any axial gradiometer sensor, the difference between the highest and lowest value of the signal during the segment exceeded 4 × 10^−12^ T. All retained data segments were concatenated into a single (sensors x time) data matrix. Prior to running the ICA, principal components analysis (PCA) was run on the data (using MNE’s ICA tools). The ICA was then conducted using the first 80 principal components. With the assistance of various diagnostic plots, components that appeared to be related to eye movements, heartbeats, and muscle activity were identified.

Data for all other analyses (apart from the ICA) were high pass filtered at 0.1 Hz (zero-phase FIR), and the previously identified artefactual ICA components were removed. Again, only data from blocks of interest were retained. Further automated routines were applied for the detection of bad sensors and bad sections of the data. Jumps were detected by convolving MEG sensor time series with a jump template, implemented using a previously developed routine (see “annotate_jumps” in the code associated with Wilming et al., 2020; detection cut off 25). Sensors with more than 80 detected jumps, plus the number of different blocks being analyzed, were marked as bad, rather than excluding temporal sections of the data. Further bad temporal sections of data were identified using MNE’s muscle detection routine (z-score threshold of 10). We also marked as bad any section of data where head position was further than 12 mm from mean head position. Finally, we detected periods affected by additional large artifacts by low-pass filtering a copy of the data at 1.5 Hz, dividing the data into overlapping 3 s segments, and marking as bad any sections in which the difference between the highest and lowest signal value on any axial gradiometer sensor exceeded 6 × 10^−12^ T.

Depending on the analysis, either this cleaned continuous data was used, or at this stage the data was divided into epochs. Analysis-specific details regarding epoching are described below. Epochs were rejected if either (a) they overlapped with sections of the data previously marked as bad, or if (b) the difference between the highest and lowest signal value on any axial gradiometer sensor exceeded 2.4×10^−11^ T. Following division into epochs, data was down sampled to 400 Hz (using MNE’s resampling routine that first applies an anti-aliasing filter), and the first and final 0.15 s were removed from epochs to minimize edge artifacts.

#### Spectral analysis

At this stage in some analyses, time-variant spectral analysis was conducted, producing complex-valued time-frequency representations of the data (i.e., Fourier coefficients that contain both amplitude and phase information). The analysis-specific details are described below. Regardless of whether or not a time-frequency representation was computed, as a next step, and prior to source reconstruction, we decimated epoched data, retaining only 1 in every 10 samples (leaving us with a sampling rate of 40 Hz for epoched, source-localized data). (Sampling of continuous data is described below.)

#### Source reconstruction

For each participant we reconstructed cortical surfaces from the participant’s MRI scan using FreeSurfer (recon-all routine; Dale et al., 1999; Fischl et al., 1999; Fischl, 2012). To define regions of interest (ROIs, see below), we aligned the HCP-MMP 1.0 parcellation (Glasser et al., 2016; Mills, 2016; Van Essen et al., 2012) to the individual cortical surface reconstruction of each participant. Additionally, we applied an fMRI-based label for the primary motor cortex (M1) hand region (de Gee et al., 2017). Alignment between MRI and MEG data was conducted on a session-by-session basis, using the average position of the three fiducial coils during all non-artifact data in localizer and main experimental blocks (not just the stimulus-locked epochs from main experimental blocks as we had preregistered; Wilming et al., 2020; MNE’s co-registration routines).

We used linearly constrained minimum variance (LCMV) beamforming (i.e., spatial filtering) to estimate neural activity at the cortical source level (Van Veen et al., 1997). The procedures for computing the data covariance matrix were specific to each analysis, and are described in detail below, but we always used data without time-frequency analysis applied for computing covariance matrices. Three-layer head models were generated with the use of FreeSurfer’s “mri_watershed” routine (Ségonne et al., 2004) and the MRI from each participant (this routine differs from the routine stated in the preregistration, because we found “mri_watershed” to lead to better head models). For three participants we found the three-layer head models to be inadequate and resorted to using a single layer (a further difference from the preregistration). These were then used (applying MNE’s default values for layer conductivities) – in conjunction with a participant-specific surface-based source space (computed using MNE’s “setup_source_space” routine) – to generate forward solutions (i.e., lead-fields). Each LCMV beamformer was computed using the relevant lead-field and covariance matrix. We used eigendecomposition to find the source orientation at each considered source location (“vertex”) on the cortical surface that maximized power. The LCMV beamformers produced in this manner, which are sets of spatial filters, were applied to the (cleaned) signal at the MEG sensors, if no time-frequency analysis had been applied. If time-frequency analysis had been applied, then each frequency was considered in turn, and the spatial filter was applied to the complex-valued representation (for the considered frequency) at the MEG sensors (Wilming et al., 2020). In the case of time-frequency analysis, the resulting source estimates were finally converted into power by taking the absolute value and squaring. Baselining was not conducted.

In sum, this procedure yielded (for each epoch) estimates of activity at each time point and vertex if time-frequency analysis had not been applied. If time-frequency analysis had been applied, it yielded (for each epoch) estimates of spectral power for each time point, frequency, and vertex.

### MEG data analysis: cortex-wide decoding of latent variables

#### Preprocessing and source-localization

Preprocessing and source localization was performed following the general MEG analysis procedure described above, with the following specific settings and choices.

All parts of this analysis were based on epoched data. For analyses time-locked to cue onset we created epochs using a window from −0.5 to 1.5 s relative to cue onset. For response-locked analyses (i.e. locked to the button press) we used the window −2.2 to 2 s relative to response. In both cases, we only used cues and responses in Inferred condition blocks.

For most of the decoding analyses we did not use time-frequency analysis to compute power at different frequencies. Instead we performed source localization using the measured signal with no time-frequency analysis applied, producing a single time-course of estimated activity for each vertex. Only for Figure 6 did we compute time-frequency representations of the data, to which we then applied source reconstruction. In this case we used the sliding-window multi-taper method (Mitra & Pesaran, 1999), with settings tailored to the frequencies that we were investigating. We used a temporal window of 0.4 s (steps of 2.5 ms). The product of the temporal window length and full frequency bandwidth was 2 (implying a full frequency bandwidth of 5 Hz). We estimated complex-valued time-frequency representations for 5Hz and 10Hz. We used the representation at 10Hz as our measure of alpha activity. Due to smoothing across frequencies these representations will reflect activity at a range of nearby frequencies (specifically, 2.5 - 7.5 Hz and 7.5 - 12.5 Hz).

LCMV beamformers were computed separately for each session, and separately depending on whether we were looking at cue-locked or response-locked epochs. Specifically, for response-locked decoding, the LCMV beamformers were constructed using covariance matrices computed using the time window −0.6 to 0 s relative to response. For cue-locked decoding we used −0.2 to 0.8 s relative to cue onset. We did not baseline this data prior to the computation of the covariance matrices.

For these analyses we only used the HCP-MMP 1.0 parcellation not the fMRI-based label for the primary motor cortex hand region.

#### Decoder training and evaluation

For the decoding we used linear support vector regression (implemented in Scikit-learn with L2 loss function; Pedregosa et al., 2011). Training was performed separately for each participant, for each of the 180 parcels defined by Glasser et al. (2016), and for each time point. We performed the decoding by pooling activity patterns across homotopic parcels from both the left and right hemisphere.

Each epoch represented one case for training or testing the decoder. The features used by the decoder to predict the target were based on the estimated source activity (or power at either 5Hz or 10Hz for Figure 6) at the specific time point under consideration, at each vertex in the parcel under consideration. Hence, the total number of features would have matched the total number of vertices in the parcel in both hemispheres, however, we reduced the number of features. Specifically, z-scoring and principle components analysis (PCA) was conducted using the training data (and not held-out test data). The *n* largest principle components were retained, such that the retained components captured 95% of the variance. Hence, the final number of features was *n*.

The targets to be predicted by the decoder were latent variables from the above-described normative model. We again computed these on a participant-by-participant basis using the fitted values for hazard rate (*Ĥ*), evidence scaling (*β̂*), and decision noise (*ν*), in the normative model. In many cases, targets of the decoding were the “residuals” of these latent behavioral variables, after regressing out the effects of other variables. That is, we used one or more variables, plus an intercept term, to predict the latent behavioral variable of interest using linear regression. We then subtracted the values predicted by the regression from the true values, leaving only “residuals”.

In decoding locked to cue-onset (e.g. Figure 4), “prior” refers to the log-prior ratio immediately before observing the time-locked cue (Ψ*_n_*), while “prior (residuals)” refers to this quantity with the effects of the log-likelihood ratio of the time-locked cue (*LLR_n_*), and of the previous cue (*LLR_n−_*_1_), regressed out. “Evidence” refers to the log-likelihood ratio (*LLR_n_*) of the time-locked cue, while “evidence (residuals)” is this quantity with the effects of the log-prior ratio before observing the time-locked cue (Ψ*_n_*), and the log-prior ratio after observing the time-locked cue (Ψ*_n_*_+1_), regressed out.

“CPP” refers to the CPP having observed the time-locked cue (*CCP_n_*; where *n* indexes the time-locked cue), and “uncert.” refers to uncertainty prior to observing the time-locked cue (*− |*Ψ*_n_|*). “CPP x LLR” refers to the interaction between CPP and LLR (i.e., *CPP_n_ × LLR_n_* where *n* indexes the time-locking cue). “Uncert. x LLR” refers to the interaction between uncertainty prior to observing the time-locked cue and LLR (i.e., *− |*Ψ*_n_| × LLR_n_*). “CPP x LLR (residuals)” and “Uncert. x LLR (residuals)” are these quantities with the effects of *LLR_n_*, and the log-prior ratio before and after observing the time-locked cue (Ψ*_n_* and Ψ*_n_*_+1_) regressed out.

In decoding locked to behavioral response (Figure 5), “posterior” refers to the log-posterior ratio (*L_n_*) after observing the most recent cue (i.e. after the last cue of the preceding ITI, which we have indexed by *n*). “Posterior (residuals)” refers to this quantity with the effects of the log-likelihood ratio of the most recent cue (*LLR_n_*), and of the second most recent cue (*LLR_n−_*_1_), regressed out. “Evidence (residuals)” refers to the log-likelihood ratio of the most recent cue (*LLR_n_*), with the effect of the log-posterior ratio after observing the most recent cue (*L_n_*), and after observing the second most recent cue (*L_n−_*_1_), regressed out.

Training and evaluation was conducted using “leave one-session out” cross-validation. That is, one of the four MEG sessions was chosen at random to be the test session. Data from the other three MEG sessions was used for training the decoders, and the decoder performance was then evaluated on the data from the held-out test session. Performance in the held-out test data was evaluated by predicting the targets of the decoding using the trained decoders, and correlating the predicted targets with the true targets (Pearson correlation coefficient). This procedure was repeated a further three times, so that each session was used once as the test session. Performance was averaged over each train/test split of the data.

This form of cross-validation provides a very stringent test: the data used to train and test the decoders were collected on separate days (and the participant was potentially positioned somewhat differently relative to the MEG device’s gradiometers). The regressions described above, to produce the “residuals” of the decoding targets, were conducted on a session-by-session basis to preclude the possibility of subtle “data leakage” across the training-test split of the data (Kapoor & Narayanan, 2023). The z-scoring and PCA of the training data can be viewed as a transformation of this data. Exactly the same transformation was applied to the test data (i.e., we subtracted off the mean of the training data, divided by the standard deviation of the training data, and used the principle components from the training data).

#### Decoder performance statistics

The approach allows us to calculate cross-validated estimates of decoder performance for each participant, cortical region, decoding target, time point of training data, and time point of evaluation data (from the held-out session). Here “time point” refers to time relative to the time-locking event. Apart from for the generalization matrix plots, we looked at the performance of the decoders evaluated on held-out data from the same time point as the time point of the data on which they were trained.

Except where explicitly stated, we performed statistics on a coarser spatial resolution than full granularity (180 parcels per hemisphere) of the anatomical atlas described above: separately for each participant, we averaged the decoding performance data in each of the 22 larger groupings of parcels also set out by Glasser et al. (2016, supplementary neuroanatomical results). In addition, we moved to a courser temporal resolution for all analyses, except for the temporal generalization analysis described below. We had trained and evaluated decoders at 0.025 s intervals. We now binned the performance data within temporal windows, and averaged the cross-validated performance within these bins, again separately for each participant. For decoding relative to cue onset we used the following windows: Precue (*−*0.05 *<*= *t <*= 0.025 s relative to cue onset), early post-cue (0.05 *<*= *t <*= 0.125 s), and late post-cue (0.15 *<*= *t <*= 0.225 s). For decoding relative to responses we used the following windows: Early pre-response (*−*1 *<*= *t <*= *−*0.5 s relative to response) and late pre-response (*−*0.5 *<*= *t <*= 0 s). The only exception is Figure 6 where power at a single time point is already computed using the neighboring time points (due to the the nature of time-frequency analysis). Here, we did not further average within temporal windows and instead used two timepoints: Early post-cue (0.1 s) and very late post-cue (0.475 s).

We only color code statistically significant (across participants) regional decoding scores on the cortical surface. We performed a one-sample T-test for each time window and region (i.e., group of parcels). We corrected the resulting p-values for the multiple comparisons across regions (within a time window) with the false discovery rate (FDR; Benjamini and Yekutieli, 2001).

For plots showing the difference between decoder performance for two different decoding targets (e.g. the subplot in Figure 4 showing the difference in prior and evidence decoding) the difference in decoder performance was computed for each participant, time-point, and parcel, prior to moving to a courser spatial and temporal resolution.

For the so-called temporal generalization matrix, we plot the decoding performance of decoders not just evaluated with data from the same time-point (relative to time-locking event) used for training the decoders, but also from all other time points (King & Dehaene, 2014). We again averaged within the 22 groups of parcels set out by Glasser et al. (2016), and moved to a courser temporal-resolution, binning the data based on both train time and evaluation time to produce bins each containing 25 data points, that were then averaged. We only plot points that were statistically significant across participants (T-test and correction with FDR for the multiple comparisons across train time-point and test time-point combinations).

#### Gaze position control analyses

We also evaluated the prior belief decoders separately for cases in which gaze position was “consistent” and “inconsistent” with the prior. For this analysis we used the version of the eye tracking data that was additionally collected with the MEG recordings at 1200 Hz. Because this version of the eye tracking data was uncalibrated, we excluded from the analysis any participant for whom, in any session, there were issues with the raw image used for eye tracking (four participants excluded).

Cue epochs were created slightly differently for this analysis. Specifically, during preprocessing we now additionally identified periods of time with missing eye tracking data (e.g. due to blinks). Epochs containing such periods were excluded. To reduce the number of excluded epochs we used a smaller time window for epoching (−0.25 to 0.45 s relative to cue onset).

For each cue in the Inferred condition we computed the average gaze position in the vertical direction (“y-gaze”) in the “pre-cue” time window (*−*0.05 *<*= *t <*= 0.025 s relative to cue onset). On a session-by-session basis we performed a median split of these average y-gaze positions, and a separate median split of the prior residuals (defined in the same way as before). We then separated cues based on whether the gaze position was consistent or inconsistent with the prior residuals. “Consistent” means that (i) y-gaze was more towards the top of the screen than the median, and the prior residual was on the side of the median split that most supported the context associated with cues on the top half of the screen, or, (ii) y-gaze was on the side of the split towards the bottom of the screen and the prior residual was on the side of the split associated with cues on the bottom half of the screen.

Decoding was performed as before, with the following key difference. Although we trained the decoders on the data for all the relevant cues, we evaluated the decoders separately on gaze consistent and gaze inconsistent cues. We again used leave-one-session out cross validation, meaning that, in each iteration of the cross-validation, the gaze consistent and gaze inconsistent cues used for evaluation all came from the held out MEG session that was not used for training the decoders.

### MEG data analysis: Correlated variability of stimulus and action codes

We performed an analogous analysis to that performed by van den Brink et al. (2023) using fMRI data, in the MEG data that we collected. Data from main experimental blocks during the MEG sessions in both the Instructed and Inferred conditions were used.

#### Preprocessing and source-localization

Preprocessing and source localization was performed following the general MEG analysis procedure described above, with the following specific settings and choices.

While some parts of the analysis used the cleaned continuous data, other parts relied on data that was further divided into epochs around specific events. For stimulus-locked epochs we used a window from −0.8 to 2 s relative to stimulus onset. For response-locked (i.e., action) epochs we used a window from −2.2 to 2 s relative to subjects’ behavioral reports (button presses).

Time-variant spectral analysis of the artifact-cleaned, sensor-level MEG data was performed using the sliding-window multi-taper method (Mitra & Pesaran, 1999). We used a temporal window of 0.25 s for the range from 4 to 9 Hz, and a temporal window of 0.1 s for the range from 10 to 150 Hz (steps of 2.5 ms). Frequency bandwidth was determined by setting the product of the temporal window length and full frequency bandwidth to 2. Step size in the frequency domain was 1 Hz for the lower frequency range and 5 Hz for the higher frequency range.

For each session, one set of spatial filters (i.e., a LCMV beamformer) was computed (for visual cortical regions of interest; ROIs) using the stimulus-locked data from the main experimental blocks, and another set of spatial filters was computed (for action-related ROIs) using the response-locked data. (ROIs described below.) Estimates of the data covariance matrix for the LCMV beamformer were computed across the time window from 0 to 1 s relative to stimulus onset for stimulus-locked epochs, and −1 to 1 s relative to button press for response-locked epochs. Differing slightly from the preregistration, we demeaned the data used for computation of the covariance matrix by, for each epoch and MEG channel, subtracting off the mean value over the included window. The thereby produced spatial filters were applied to the complex-valued time-frequency representation of the (continuous or epoched) sensor-level data (i.e., it was applied to the Fourier coefficients).

We considered a set of visual cortical regions and a set of action-related ROIs. As described below, ROIs were only included in the final correlated variability analysis if decoding using those regions achieved a certain level of performance. The considered visual regions were (a) V1, (b) early visual cortex, (c) dorsal stream visual cortex, (d) ventral stream visual cortex, (e) MT+ complex and neighboring visual areas (LOC complex), all as defined by Glasser et al. (2016, supplementary information). The considered action-related frontal cortical regions were (a) the M1-hand region from de Gee et al. (2017), and the following clusters of Glasser et al. (2016) parcels: (b) 6a and 6d, (c) FEF and PEF, and (d) 6v and 6r. Overlap between parcels of the Glasser et al. (2016) atlas and the M1 label was dealt with by removing all overlapping points from the corresponding Glasser et al. (2016) parcels in somatomotor cortex.

#### Stimulus and action decoder training

We first trained stimulus and action decoders on epoched data, and we then applied the trained decoders to the preprocessed continuous MEG time series. We used linear support vector classifiers (implemented in Scikit-learn; Pedregosa et al., 2011). Considering first the training, for each participant we trained one stimulus decoder (horizontal vs. vertical grating) per visual ROI, and one action decoder (left vs. right button press) per motor ROI. We trained using all main experimental task data from all sessions. Because decoder performance can be affected by unbalanced datasets, epochs were dropped where necessary to ensure equal numbers of epochs for the two stimuli, and equal numbers of epochs for the two actions (performed on a participant-by-participant basis).

Source activity estimates were produced for each epoch, as described above. The features given to the stimulus decoders to estimate stimulus orientation were based on the power at each vertex in the left- and right-hemispheric portion of each ROI, and at each considered frequency (implying a total number of features equal to the number of frequencies multiplied by the number of vertices in the ROI across both hemispheres). However, the number of features was reduced prior to training the decoder by z-scoring the data, and then retaining only the first n largest principal components, where n is set such that the retained components captured 95% of the original variance. Z-scoring and principal components analysis (PCA) were conducted on the training data. As training data, we used the values of the features at every time point between 0.200 and 0.275 s relative to stimulus onset, for every epoch (implying a total number of samples for the decoder to train on equal to the number of time points in the considered interval multiplied by the number of epochs). The action decoders were trained in the same manner. As samples for the action decoders, we used the value of the features at every time point between −1.50 and −0.125 s relative to the action, for every epoch. The specific time windows were selected and preregistered based on pilot results, which showed robust decodability of stimulus orientation and chosen action, respectively, using these intervals.

Note that the features at different time points within the same epoch are likely to be highly correlated (due to overlap of the spectral estimation windows, as well as genuine autocorrelation of neural activity), and hence may provide largely redundant samples for training. We nevertheless include multiple time points from the same epoch so that the decoders learn to classify reliably based on largely time-invariant patterns in the data, not just at particular time points of the evoked responses. This is because in our correlated variability analysis, the decoders are applied to ongoing activity measured during the intertrial intervals (ITIs): Decoders that only work at specific time points of evoked responses will not be suitable.

#### Exclusion of regions of interest based on decodability of evoked responses

Our correlated variability analysis hinged on the ability to track ongoing fluctuations of stimulus and action-selective cortical activity in different ROIs via the decoders trained on the ROIs’ stimulus- and action-evoked responses. A prerequisite for doing so is obtaining robust stimulus and action decoding from those evoked responses, for all ROIs included. If this criterion is not met, the corresponding ROIs thus need to be excluded from the analysis. To this end, we used the following procedure.

For each sensory (visual cortex) and action (frontal cortex) ROI, we excluded that region (and the corresponding decoder) from the remainder of the correlated variability analysis, if a certain level of decoding performance was not achieved. To evaluate decoding performance in an ROI we looked at the four-fold cross-validated accuracy of decoders (stimulus or action, as appropriate) trained in the same manner as usual (apart from the use of cross-validation). We produced a single accuracy value per participant and region by computing, for each fold, average accuracy on the held-out data (but only using time points between 0.200 and 0.275 s relative to stimulus onset, and time points between −0.200 and −0.125 s relative to actions), and then averaging across folds.

We compared the average accuracy values (one for each participant) to 50% (i.e., chance) using a permutation t-test (10,000 permutations, each iteration involved a 0.5 probability of flipping each original datapoint around the reference value). If a given ROI did not achieve a statistically significant level of decoding accuracy (at a p-value of 0.05, one-sided test), that ROI was excluded from the remainder of the analyses.

#### Application of stimulus and action decoders to continuous data

The stimulus decoders, constructed as described above, were applied to the continuous time series of all decoding features per visual ROI, yielding a single (scalar) ROI time-series of continuous decoder output throughout each task block for each ROI (performed on a segment-by-segment basis, described below). The same process was conducted with the action decoders and the action-related ROIs. To this end, the continuous pre-processed data was first down-sampled to 400Hz, as was previously done to the epoched data.

Consider the computation of stimulus decoder output for the measurement from a particular participant, session, and visual cortical ROI. The sensor-level time-frequency representation was computed as before (but this time with no subsequent decimation of the data), source-level estimates of power were obtained (using the participant and session specific beamformer that was constructed by using the covariance matrix from the stimulus epochs), this data was then z-transformed by subtraction of the mean of training data and division by the training data standard deviation. The result was projected onto the principal components previously computed from the training data (see above). The (participant specific) stimulus decoder for this visual ROI was applied to the resulting features. (The z-transform and principal components from the training data were used to ensure correspondence of the decoding features in decoder training and application of decoders to continuous data.) For producing the continuous action decoder output for an action-related ROI, the beamformer computed from the covariance matrix in the action epochs, and the action decoder for that region was used instead.

Artifacts in the continuous data were dealt with in the following manner. We moved through the continuous data with a step size of 1 second. At each step, the data around the corresponding timepoint was evaluated for inclusion/exclusion by looking at a small window around the timepoint (0.5 s prior to 1.5 s following). If any part of the data within that window was marked as bad during the pre-processing then the associated data was not included. On the other hand, if no data within the window was marked as bad, then this segment had the relevant beamformer and decoder applied, as described, before being trimmed to 0.1s prior to 1.1s following the timepoint under consideration. All included segments were recombined (respecting excluded data segments by inserting “not a number” values in the corresponding positions) to produce continuous, scalar time series of stimulus and action decoder outputs.

#### Removal of evoked responses

Prior to conducting the main correlation analysis between stimulus decoder and action decoder output time courses, we regressed out stimulus-locked and response-locked evoked responses in these time courses. That is, separately for each stimulus decoder and each action decoder output (i.e., separately for each ROI), and separately for each participant, (but simultaneously for the data from all sessions) we ran a regression of the following form,

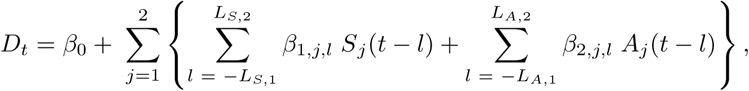

where *D_t_* is the value of the decoder output (stimulus or action) for the current ROI at time step *t*, the *β* values are the various regression coefficients (and are unrelated to the *β* parameter of the normative model), *j* indexes the specific stimulus or action, and *l* indexes time-lag. That is, we considered a range of lags from before the time-locking event, to well after the time-locking event (see below), and estimated at each lag the effect of stimuli and actions. *L_S,_*_1_ and *L_S,_*_2_ set the limits of the lags considered in the regression for the effects of the stimulus, while *L_A,_*_1_ and *L_A,_*_2_ play the same role for the effects of the action. We set these limits such that the number of lags considered covers the period −1 to 8 s relative to stimuli, and −8 to 8 s relative to action. *S_j_*(*k*) is 1 if there was an onset of the specific stimulus *j* (horizontal or vertical) at time step k, and it is 0 otherwise. Similarly, *A_j_*(*k*) is 1 if there was an onset of the specific action *j* (left or right) at time step *k*, and is 0 otherwise.

After the described linear regression model had been fit, the fitted coefficients were used to produce an estimated decoder output time series, 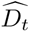, which was then subtracted from the original decoder output time series. This produced a decoder output time series with stimulus and action-locked effects regressed out. Note that for both the stimulus and action decoder output time courses we regressed out both stimulus-locked and action-locked effects.

#### Decoder output correlations

For this analysis we used the prepared decoder output time series with the stimulus- and action-locked effects removed (with the total number of time courses matching the total number of included ROIs). In this analysis we wanted to examine the effects of ongoing fluctuations in stimulus and action-encoding patterns, not stimulus- and action-evoked responses. Therefore, we only included time points occurring during the inter-trial-interval (ITI), and specifically only those time points that were at least 0.8 s following the action in the previous trial, and at least 0.125 s before onset of the upcoming grating stimulus. We excluded time points within a window of −1.325 to 0.8 s relative to any other detected button presses.

The lateralization of beta-band power has been observed to increase prior to an action (decreased contralateral vs. ipsilateral to response hand), and to flip following an action (increased contralateral vs. ipsilateral; Pfurtscheller and Lopes da Silva, 1999; Urai and Donner, 2022). This “beta rebound” has been observed to last longer than the 0.8 s exclusion window that we use. This activity is not an issue for our analyses because (a) we regressed out evoked activity in decoder output time courses using a large window around actions (see above), and (b) any residual activity will harm decoder performance, rather than leading us to draw false conclusions: Action decoders were trained on a period prior to the action when beta lateralization is expected to be in the opposite direction to the beta rebound, and therefore any residual effects of the beta rebound will work to reduce the performance of these decoders.

We were interested in the specific structure of correlations (described below) between stimulus decoder outputs and action decoder outputs. However, we have multiple stimulus decoder output series (one for each visual ROI) and multiple action decoder output series (one for each motor ROI). We first computed the full (Pearson) correlation matrix of the decoder outputs of all pairs of visual and action-related ROIs. We then averaged these correlation values selectively for the (unique) visual-action pairs (to which our predictions pertain) as well as the (unique) visual-visual and action-action pairs (not including a pair formed of an ROI and itself; for control; see corresponding matrices in van den Brink et al., 2023).

This procedure was conducted separately for data from the Instructed condition, and for data from the Inferred condition. For the Instructed condition, we computed the above correlation matrices for timepoints when the stimulus-response mapping context – provided unambiguously to participants through noiseless cues – was “Context 1”, and a second average correlation for timepoints where the context was “Context 2”. A conceptually analogous, but more complicated, division is appropriate for the Inferred condition. This analysis was based on latent variables computed (on a participant-by-participant basis) using the normative behavioral model with fitted hazard rate (*H*), evidence scaling (*β*), and decision noise (*ν*; described above). In particular, we looked at the estimated belief of participants regarding the active context. We computed two correlation matrices for the Inferred condition, one for the timepoints at which the estimated belief of a participant supported “Context 1”, and one for the timepoints at which the estimated belief of a participant supported “Context 2”. For each participant, we finally computed four mean correlation values between stimulus-action decoder outputs (i.e., collapsed across all visual-action pairs): One correlation for each context (or context belief) within each of the two conditions (Instructed vs. Inferred).

#### Tests

This procedure leaves us with four stimulus-action correlation values per participant. Our two main tests in the correlated variability analysis were conducted on these values. We predicted that fluctuations in stimulus- and motor-encoding patterns would covary in a way (specifically, with a sign) that tracks the active (or Inferred) context (van den Brink et al., 2023). All correlations were computed such that positive values indicate that horizontal grating stimulus-encoding patterns are associated with left action-encoding patterns, i.e., that correlations in intrinsic fluctuations are consistent with Context 1. Hence, the correlation values produced using data points when “Context 1” is active (Instructed condition), or believed to be active (Inferred), should be more positive than those produced when “Context 2” is active or supported.

In the Instructed condition our main test was conducted on the differences between the correlation values when “Context 1” was active and those when “Context 2” was active (one difference value per participant). We used a one-sided permutation test to compare these values to zero. We predicted that the values would be more positive under “Context 1” (preregistered).

In the Inferred condition, our main test was analogous: Here we performed a one-sided permutation test, comparing to zero the differences between the correlation values when estimated belief supported “Context 1” and the correlation values when estimated belief supported “Context 2”. Again, we expected the correlation values would be more positive under “Context 1” (preregistered).

We conducted four further follow-up tests on the average stimulus-action correlations, which were directly motivated by our hypotheses (preregistered). In the Instructed condition, we predicted that the correlation values when “Context 1” was active would be positive. This was tested with a one-sided permutation test comparing the values across participants to zero. An analogous test was conducted for the prediction that correlation values when “Context 2” was active would be negative. In the same manner, in the Inferred condition we tested whether the correlation values were positive when “Context 1” was supported by Inferred belief, and negative when “Context 2” was supported.

We also conducted control tests on the average correlation between stimulus-stimulus decoder pairs and action-action decoder pairs (preregistered). Looking at these pairs is not informative for our main hypotheses, but they can provide a check that the decoders functioned as expected. Specifically, the output of decoders trained to decode the same target (stimulus or action) should be positively correlated, regardless of whether “Context 1” or “Context 2” is currently active. Hence, we predicted positive correlation values for both stimulus-stimulus and action-action decoder pairs, in both Instructed and Inferred conditions, and under both “Context 1” and “Context 2” (8 tests in total; each a one-sided permutation test comparing the values across participants to zero).

Permutation tests were conducted by computing a null distribution for the t-statistic using 10,000 permutations or every possible permutation, whichever number was smaller. For one-sample tests comparing a set of values to a reference value, each iteration involves a 0.5 probability of flipping each original datapoint around the reference value. We applied a p-value of 0.05 as the threshold for statistical significance. No corrections for multiple comparisons were made because the tests were specified a priori through preregistration.

#### Preregistration

The correlated variability analyses were preregistered (https://osf.io/h4×5s). This preregistration was separate to, and later than, the pupil preregistration. While behavioral modeling and pupil analyses had progressed since the previous preregistration, the collected MEG data that we focus on in the correlated variability analysis had not been analyzed yet. The conducted analysis differed in some minor ways from the preregistered analysis. These differences are noted in the above description.

## Acknowledgments

We are grateful to everyone who participated in our study, and to Karin Reimann, Christiane Reissmann, Roger Zimmermann, Barbora Schwarzová, Marlene Petersson, Annika Wermuth, and Gina Monov for collecting data with us, or directly supporting our data collection. We thank Peter Murphy for support in setting up MEG source localization, and Joshua Gold for providing feedback on an earlier version of the results.

The HCP-MMP 1.0 parcellation, which we used, was developed using data from the Human Connectome Project, WU-Minn Consortium (Principal Investigators: David Van Essen and Kamil Ugurbil; 1U54MH091657) funded by the 16 NIH Institutes and Centers that support the NIH Blueprint for Neuroscience Research; and by the McDonnell Center for Systems Neuroscience at Washington University.

## Data availability

All collected data, apart from the MRI scans, will be made publicly available upon publication. MRI scans will not be included because they constitute personally identifiable data.

## Code availability

All custom code written for the study will be made available on publication.

## Author note

Large parts of the text in the “Methods” section are taken from our two preregistrations.

## Supplementary figures

**Figure 2 – Supplemental Figure 1:**
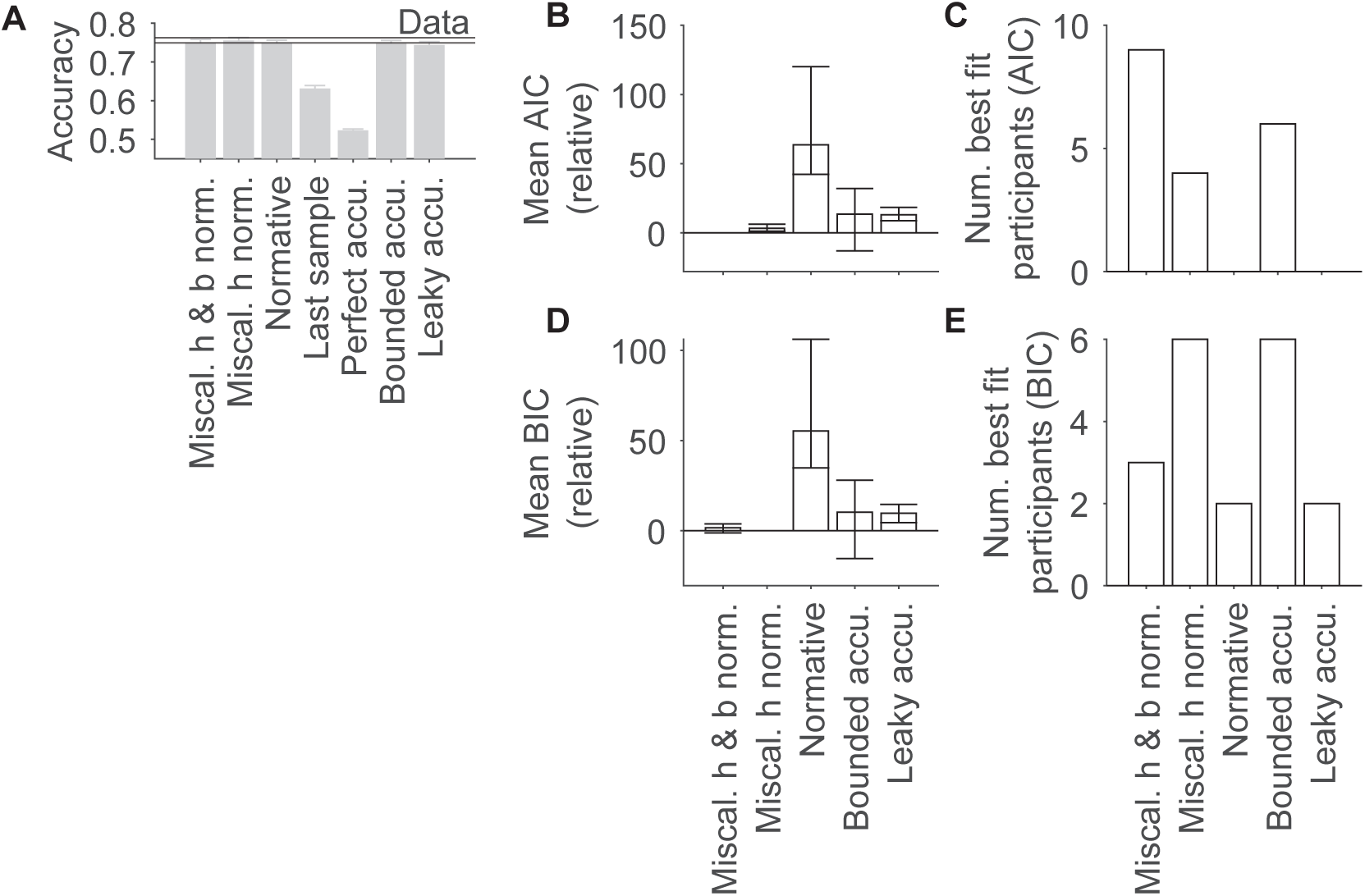
Behavioral model comparison (Inferred condition). (A) Average accuracy in the Inferred condition across participants (solid lines represent *±*1 standard error of the mean; SEM) and in the model fits (gray bars and error bars, which give +1 SEM across simulated participants). (B-E) Comparison of the goodness of fit of all models with plausible performance in (A) using AIC (B and C) and BIC (D and E). (B, D) Mean difference in AIC and BIC from the best fitting model and 95% confidence intervals on these differences. (C, E) The number of participants for which each model was the best fitting model. Further details in Methods.

**Table 2:** Secondary preregistered statistical tests of the correlated variability analysis. These tests were conducted on the correlations between decoder outputs and are described in the Methods. Figure 8 displays the associated data.

| Figure | Condition | Decoders | Active context | Sample size | T-statistic | p-value |
| --- | --- | --- | --- | --- | --- | --- |
| Figure 8A | vis-act | Instructed | 1 | 19 | -1.48 | 0.923 |
| Figure 8A | vis-act | Instructed | 2 | 19 | -0.324 | 0.621 |
| Figure 8D | vis-act | Inferred | 1 | 19 | -1.35 | 0.906 |
| Figure 8D | vis-act | Inferred | 2 | 19 | -2.26 | 0.983 |
| Figure 8B | vis-vis | Instructed | 1 | 19 | 29.6 | $1.00 \times 10^{-4}$ |
| Figure 8B | vis-vis | Instructed | 2 | 19 | 33.7 | $1.00 \times 10^{-4}$ |
| Figure 8C | act-act | Instructed | 1 | 19 | 25.9 | $2.00 \times 10^{-4}$ |
| Figure 8C | act-act | Instructed | 2 | 19 | 25.2 | $1.00 \times 10^{-4}$ |
| Figure 8E | vis-vis | Inferred | 1 | 19 | 29.1 | $1.00 \times 10^{-4}$ |
| Figure 8E | vis-vis | Inferred | 2 | 19 | 32.6 | $1.00 \times 10^{-4}$ |
| Figure 8F | act-act | Inferred | 1 | 19 | 21.9 | $1.00 \times 10^{-4}$ |
| Figure 8F | act-act | Inferred | 2 | 19 | 22.3 | $1.00 \times 10^{-4}$ |

**Figure 2 – Supplemental Figure 2:**
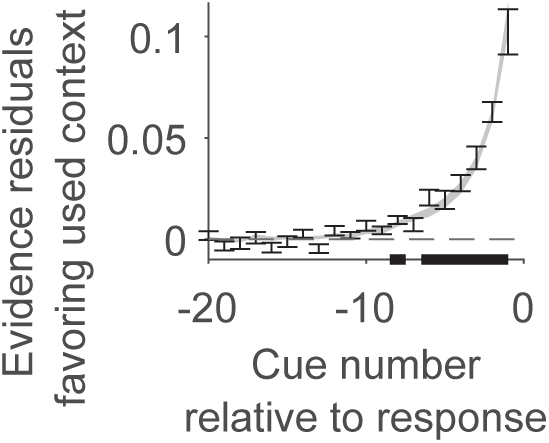
Alternative quantification of evidence weighting for higher-level decision in the Inferred condition. Average evidence residuals favoring the context used, both in the participants (error bars), and in the miscalibrated h and b normative model (shading). Evidence residuals are a measure of evidence that are not correlated between time-points. Average evidence residuals signed, such that positive values indicate support for the used context, will be positive at time-points used to determine the context, and zero at time-points that do not influence the choice (see Methods; Resulaj et al., 2009). Error bars and shading give *±*1 standard error of the mean. Bold lines on the axis represent periods in which the participant data differed significantly from zero (two-sided permutation cluster-based T-test with threshold-free cluster enhancement, *p <*= 0.05).

**Figure 2 – Supplemental Figure 3:**
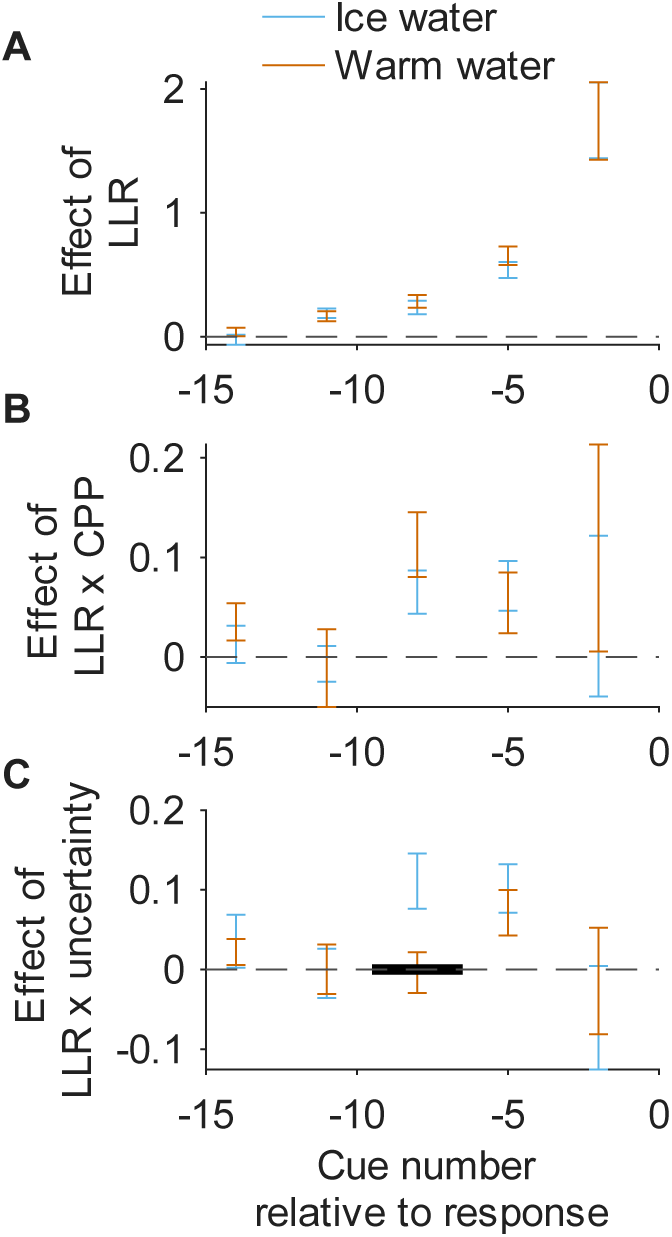
The regression analysis from Figure 2, performed on the real data, but additionally examining effects of the cold pressor test. This regression analysis was conducted in the same way as for the main figure, except additional terms were included in the regression for the use of the cold pressor test (“ice water”; see Methods). This allowed us to plot effects with the cold pressor test (“ice water”) and without (“warm water”). Error bars correspond to *±*1 standard error of the mean across participants. Unlike in the main figure, we assessed for significant differences between the “ice water” and “warm water” conditions, indicated by bold lines on the x- axis (*p <*= 0.05; two-sided; permutation cluster-based T-test with threshold-free cluster enhancement; see Methods). Results were qualitatively and quantitatively similar under the two conditions. Only for the effect of LLR modulated by uncertainty was there a significant difference between the conditions.

**Figure 3 – Supplemental Figure 1:**
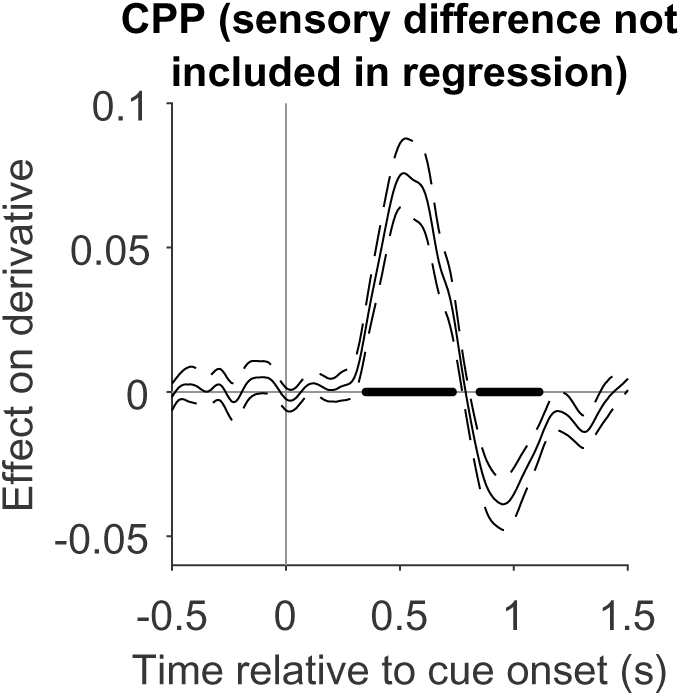
The effect of CPP on the derivative of pupil diameter when not including a term for sensory difference (preregistered). This variant of the CPP analysis presented in Figure 3 was conducted to test the robustness of that result. Dashed lines represent *±*1 standard error of the mean. Bold lines on the axis represent periods in which the data differed significantly from zero (non-parametric permutation test, with false discovery rate correction, two-sided, *q <*= 0.05). Further details in Methods.

**Figure 4 – Supplemental Figure 1:**
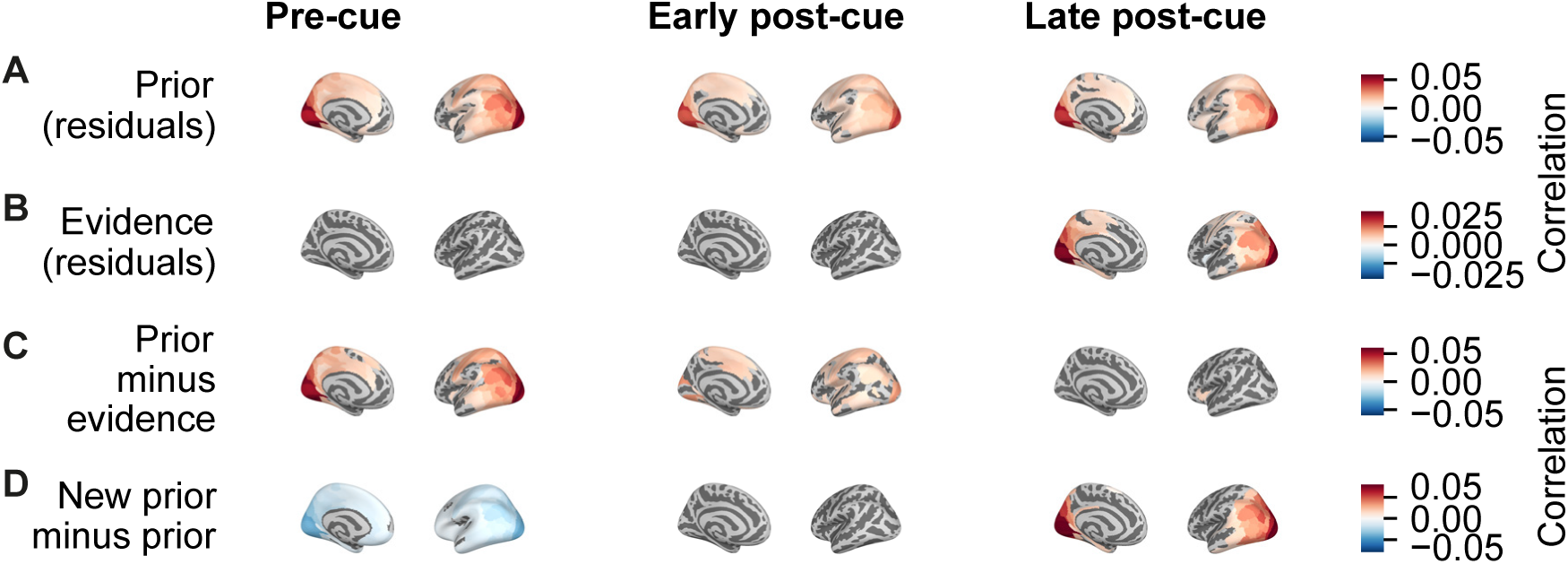
Cortical distribution of decoding performance using a fine-grained cortical parcellation (180 areas per hemisphere) as defined by Glasser et al. (2016). Beyond this, the analysis and plotting is identical to that used in the corresponding subplots of Figure 4. Decoding is again performed based on activity patterns in homotopic areas from both hemispheres, and decoding performance is displayed on the left hemisphere for visualization. Only regions with decoding significantly different from zero are shaded (assessed using two-sided one-sample T-tests corrected using false discovery rate; *q <*= 0.05).

**Figure 4 – Supplemental Figure 2:**
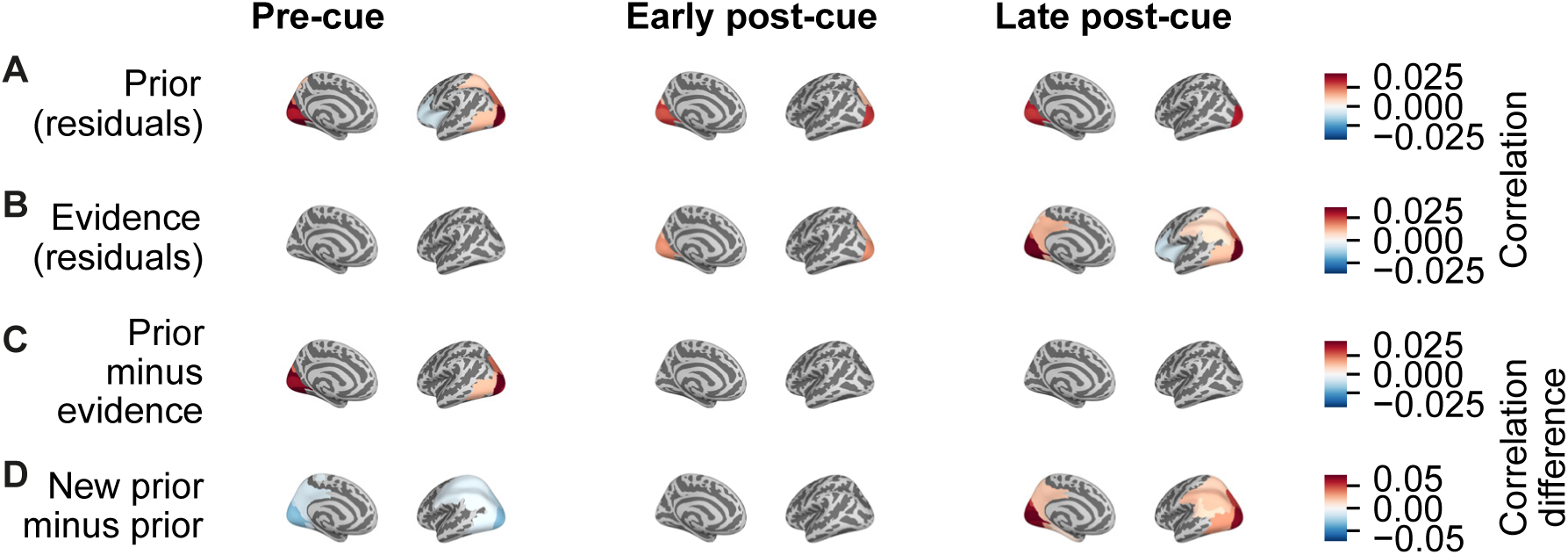
Cortical distribution of decoding performance, relative to the decoding performance in primary auditory cortex (A1). Analysis and plotting is identical to that used in the corresponding subplots of Figure 4 except that we computed, separately for each participant, time-point and cortical region, the difference in the decoding performance (measured using correlation) between the region under consideration, and the decoding performance in the A1 parcel. Hence, the figure shows differences in decoder performance from the performance achieved in A1. Only regions with decoding (differences) significantly different from zero are shaded (assessed using two-sided one-sample T-tests corrected using false discovery rate; *q <*= 0.05)

**Figure 4 – Supplemental Figure 3:**
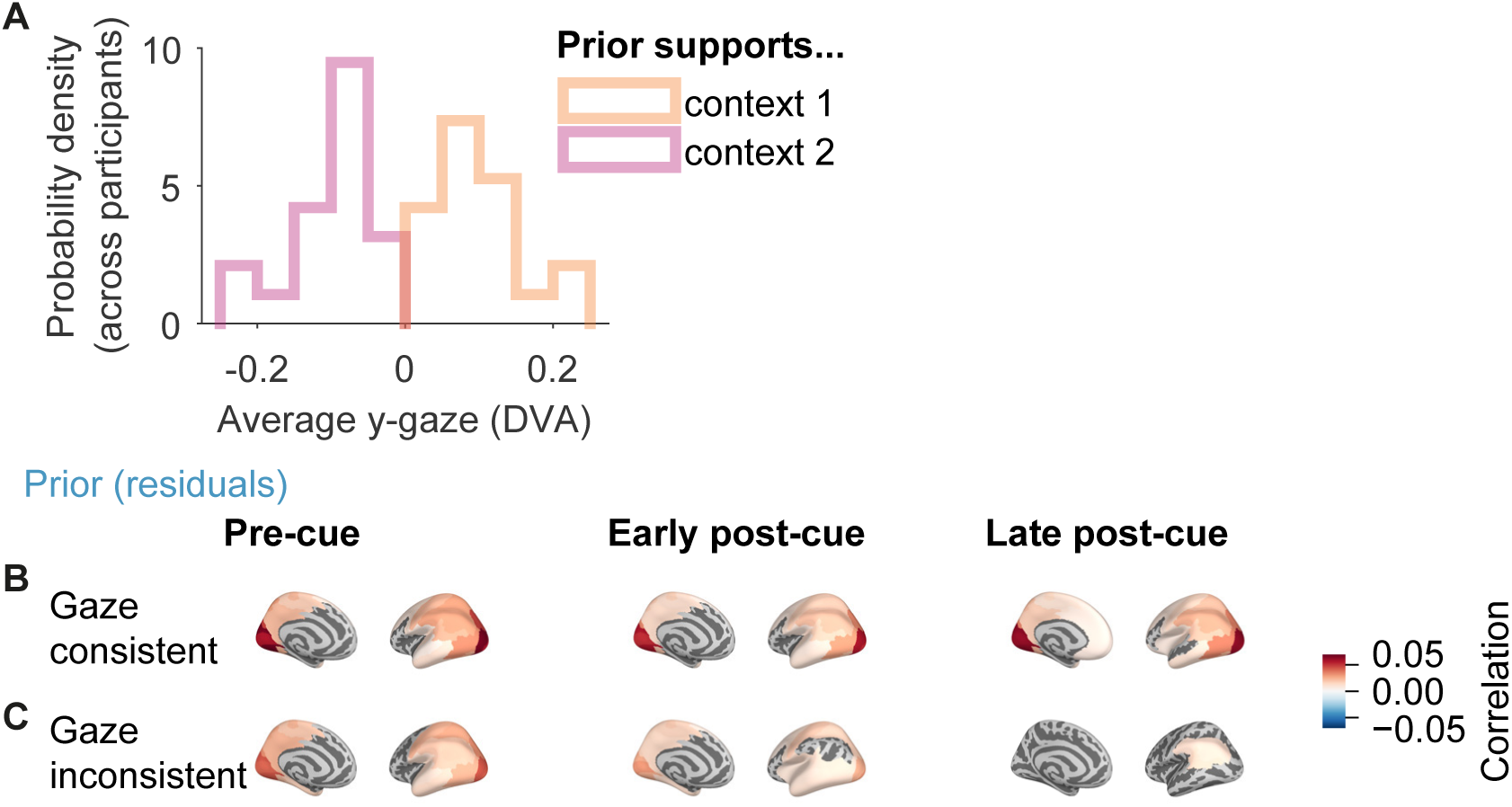
Average y-gaze position and performance of prior belief decoders trained on all Inferred condition data but evaluated contingent on gaze position. (A) Separately for each participant we computed average gaze position along the y-axis of the screen (i.e. along the vertical axis) in degrees of visual angle (DVA). This was performed separately depending on the context supported by the prior, using Inferred condition data from the pre-cue time window. The histogram represents the distribution across participants. (B, C) Decoding was performed as in main Figure 4, with two exceptions. First, we excluded participants for whom there had been issues with the raw eye tracking image (four excluded; see Methods). Second, cross-validated performance was evaluated separately on two types of cue, “gaze-consistent” cues and “gaze inconsistent” cues, based on median splits of both y-gaze position (during pre-cue time window) and the prior residuals. “Gaze consistent” refers to cues for which y-gaze position and the (residual) prior were on consistent sides of the median split. E.g. gaze was in the upper half of the visual field, and prior was on the side of the median split that was associated with cues in the upper-half of the visual field. “Gaze inconsistent” refers to cues for which y-gaze position and the (residual) prior were on opposite sides of the median splits. Maps of cortical distribution for (B) gaze-consistent priors and (C) gaze-inconsistent priors. Only regions with decoding significantly different from zero are shaded (assessed using two-sided one-sample T-tests corrected using false discovery rate; *q <*= 0.05). Further details in Methods.

**Figure 4 – Supplemental Figure 4:**
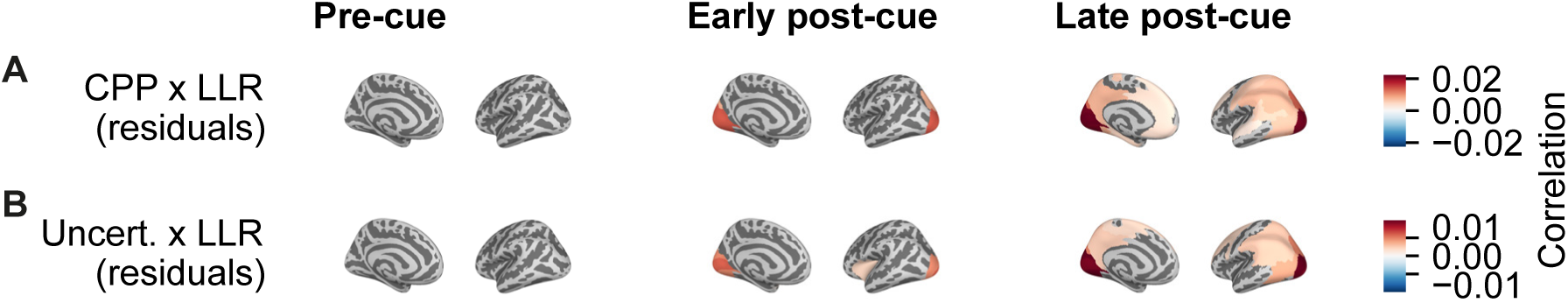
Modulation of evidence decoding by CPP or uncertainty. Decoding was performed separately for different brain regions using source activity estimates computed from measured MEG signals, and cross-validated performance was evaluated in the same way as before. (A) Decoding of the interaction between the CPP and the LLR associated with the cue, with effects of LLR and log-prior ratio regressed out. (B) Decoding of the interaction between uncertainty before observing the cue and LLR, with the effects of LLR and log-prior ratio regressed out. Further details in Methods. Only regions with decoding significantly different from zero are shaded (assessed using two-sided one-sample T-tests corrected using false discovery rate; *q <*= 0.05).

## References

Acerbi, L., Dokka, K., Angelaki, D. E., & Ma, W. J. (2018). Bayesian comparison of explicit and implicit causal inference strategies in multisensory heading perception. PLOS Computational Biology, 14(7). 10.1371/journal.pcbi.1006110

Aston-Jones, G., & Cohen, J. D. (2005). An integrative theory of locus coeruleus-norepinephrine function: Adaptive gain and optimal performance. Annual Review of Neuroscience, 28(1), 403–450. 10.1146/annurev.neuro.28.061604.135709

Baillet, S. (2017). Magnetoencephalography for brain electrophysiology and imaging. Nature Neuroscience, 20(3), 327–339. 10.1038/nn.4504

Bastos, A. M., Vezoli, J., Bosman, C. A., Schoffelen, J.-M., Oostenveld, R., Dowdall, J. R., De Weerd, P., Kennedy, H., & Fries, P. (2015). Visual areas exert feedforward and feedback influences through distinct frequency channels. Neuron, 85(2), 390–401. 10.1016/j.neuron.2014.12.018

Benjamini, Y., & Yekutieli, D. (2001). The control of the false discovery rate in multiple testing under dependency. The Annals of Statistics, 29(4), 1165–1188. Retrieved 2024, from https://www.jstor.org/stable/2674075

Bishop, C. M. (2006). Pattern recognition and machine learning. Springer.

Bogacz, R., Brown, E., Moehlis, J., Holmes, P., & Cohen, J. D. (2006). The physics of optimal decision making: A formal analysis of models of performance in two-alternative forced-choice tasks. Psychological Review, 113(4), 700–765. 10.1037/0033-295X.113.4.700

Bouret, S., & Sara, S. J. (2005). Network reset: A simplified overarching theory of locus coeruleus noradrenaline function. Trends in Neurosciences, 28(11), 574–582. 10.1016/j.tins.2005.09.002

Brainard, D. H. (1997). The psychophysics toolbox. Spatial Vision, 10(4), 433–436. 10.1163/156856897X00357

Brink, R. L. v. d., Hagena, K., Wilming, N., Murphy, P. R., Calder-Travis, J., Finsterbusch, J., Büchel, C., & Donner, T. H. (2024). Dynamics of brainstem arousal systems and pupil size predict cortical interactions for flexible decision-making. 10.1101/2023.12.05.570327

Charles, L., & Yeung, N. (2019). Dynamic sources of evidence supporting confidence judgments and error detection. Journal of Experimental Psychology: Human Perception and Performance, 45(1), 39–52. 10.1037/xhp0000583

Chen, S., Liu, Y., Wang, Z. A., Colonell, J., Liu, L. D., Hou, H., Tien, N.-W., Wang, T., Harris, T., Druckmann, S., Li, N., & Svoboda, K. (2024). Brain-wide neural activity underlying memory-guided movement. Cell, 187 (3), 676–691.e16. 10.1016/j.cell.2023.12.035

Cohen, J. (1988). Statistical power analysis for the behavioral sciences (2nd ed.). Erlbaum Associates.

Dale, A. M., Fischl, B., & Sereno, M. I. (1999). Cortical surface-based analysis: I. segmentation and surface reconstruction. NeuroImage, 9(2), 179–194. 10.1006/nimg.1998.0395

Dayan, P., & Yu, A. J. (2006). Phasic norepinephrine: A neural interrupt signal for unexpected events. Network: Computation in Neural Systems, 17 (4), 335–350. 10.1080/09548980601004024

de Gee, J. W., Colizoli, O., Kloosterman, N. A., Knapen, T., Nieuwenhuis, S., & Donner, T. H. (2017). Dynamic modulation of decision biases by brainstem arousal systems. eLife, 6. 10.7554/eLife.23232

Ding, L., & Gold, J. I. (2013). The basal ganglia’s contributions to perceptual decision making. Neuron, 79(4), 640–649. 10.1016/j.neuron.2013.07.042

Donner, T. H., Siegel, M., Fries, P., & Engel, A. K. (2009). Buildup of choice-predictive activity in human motor cortex during perceptual decision making. Current Biology, 19(18), 1581–1585. 10.1016/j.cub.2009.07.066

Filipowicz, A. L., Glaze, C. M., Kable, J. W., & Gold, J. I. (2020). Pupil diameter encodes the idiosyncratic, cognitive complexity of belief updating. eLife, 9. 10.7554/eLife.57872

Findling, C., Hubert, F., Acerbi, L., Benson, B., Benson, J., Birman, D., Bonacchi, N., Buchanan, E. K., Bruijns, S., Carandini, M., Catarino, J. A., Chapuis, G. A., Churchland, A. K., Dan, Y., Davatolhagh, F., DeWitt, E. E. J., Engel, T. A., Fabbri, M., Faulkner, M. A., … Pouget, A. (2025). Brain-wide representations of prior information in mouse decision-making. Nature, 645(8079), 192–200. 10.1038/s41586-025-09226-1

Fischl, B., Sereno, M. I., & Dale, A. M. (1999). Cortical surface-based analysis. II: Inflation, flattening, and a surface-based coordinate system. NeuroImage, 9(2), 195–207. 10.1006/nimg.1998.0396

Fischl, B. (2012). FreeSurfer. NeuroImage, 62(2), 774–781. 10.1016/j.neuroimage.2012.01.021

Friston, K. (2010). The free-energy principle: A unified brain theory? Nature Reviews Neuroscience, 11(2), 127–138. 10.1038/nrn2787

Friston, K., & Kiebel, S. (2009). Predictive coding under the free-energy principle. Philosophical Transactions of the Royal Society B: Biological Sciences, 364(1521), 1211–1221. 10.1098/rstb.2008.0300

Fusi, S., Asaad, W. F., Miller, E. K., & Wang, X.-J. (2007). A neural circuit model of flexible sensorimotor mapping: Learning and forgetting on multiple timescales. Neuron, 54(2), 319–333. 10.1016/j.neuron.2007.03.017

Glasser, M. F., Coalson, T. S., Robinson, E. C., Hacker, C. D., Harwell, J., Yacoub, E., Ugurbil, K., Andersson, J., Beckmann, C. F., Jenkinson, M., Smith, S. M., & Van Essen, D. C. (2016). A multi-modal parcellation of human cerebral cortex. Nature, 536(7615), 171–178. 10.1038/nature18933

Glaze, C. M., Kable, J. W., & Gold, J. I. (2015). Normative evidence accumulation in unpredictable environments. eLife, 4, e08825. 10.7554/eLife.08825

Gold, J. I., & Shadlen, M. N. (2000). Representation of a perceptual decision in developing oculomotor commands. Nature, 404(6776), 390–394. 10.1038/35006062

Gold, J. I., & Shadlen, M. N. (2001). Neural computations that underlie decisions about sensory stimuli. Trends in Cognitive Sciences, 5(1), 10–16. 10.1016/S1364-6613(00)01567-9

Gold, J. I., & Shadlen, M. N. (2007). The neural basis of decision making. Annual Review of Neuroscience, 30(1), 535–574. 10.1146/annurev.neuro.29.051605.113038

Gramfort, A., Luessi, M., Larson, E., Engemann, D., Strohmeier, D., Brodbeck, C., Goj, R., Jas, M., Brooks, T., Parkkonen, L., & Hämäläinen, M. (2013). MEG and EEG data analysis with MNE-python. Frontiers in Neuroscience, 7. 10.3389/fnins.2013.00267

Gramfort, A., Luessi, M., Larson, E., Engemann, D. A., Strohmeier, D., Brodbeck, C., Parkkonen, L., & Hämäläinen, M. S. (2014). MNE software for processing MEG and EEG data. NeuroImage, 86, 446–460. 10.1016/j.neuroimage.2013.10.027

Hanks, T. D., Kopec, C. D., Brunton, B. W., Duan, C. A., Erlich, J. C., & Brody, C. D. (2015). Distinct relationships of parietal and prefrontal cortices to evidence accumulation. Nature, 520(7546), 220–223. 10.1038/nature14066

Hines, E. A., & Brown, G. E. (1936). The cold pressor test for measuring the reactibility of the blood pressure: Data concerning 571 normal and hypertensive subjects. American Heart Journal, 11(1), 1–9. 10.1016/S0002-8703(36)90370-8

Jardri, R., & Denève, S. (2013). Circular inferences in schizophrenia. Brain, 136(11), 3227–3241. 10.1093/brain/awt257

Jonikaitis, D., Xia, R., & Moore, T. (2025). Robust encoding of stimulus–response mapping by neurons in visual cortex. Proceedings of the National Academy of Sciences, 122(9), e2408079122. 10.1073/pnas.2408079122

Jordan, R., & Keller, G. B. (2023). The locus coeruleus broadcasts prediction errors across the cortex to promote sensorimotor plasticity. eLife, 12, RP85111. 10.7554/eLife.85111

Joshi, S., & Gold, J. I. (2020). Pupil size as a window on neural substrates of cognition. Trends in cognitive sciences, 24(6), 466–480. 10.1016/j.tics.2020.03.005

Kapoor, S., & Narayanan, A. (2023). Leakage and the reproducibility crisis in machine-learning-based science. Patterns, 4(9), 100804. 10.1016/j.patter.2023.100804

Khilkevich, A., Lohse, M., Low, R., Orsolic, I., Bozic, T., Windmill, P., & Mrsic-Flogel, T. D. (2024). Brain-wide dynamics linking sensation to action during decision-making. Nature, 634(8035), 890–900. 10.1038/s41586-024-07908-w

Kiani, R., Cueva, C. J., Reppas, J. B., & Newsome, W. T. (2014). Dynamics of neural population responses in prefrontal cortex indicate changes of mind on single trials. Current Biology, 24(13), 1542–1547. 10.1016/j.cub.2014.05.049

King, J.-R., & Dehaene, S. (2014). Characterizing the dynamics of mental representations: The temporal generalization method. Trends in Cognitive Sciences, 18(4), 203–210. 10.1016/j.tics.2014.01.002

Kleiner, M., Brainard, D., & Pelli, D. (2007). What’s new in psychtoolbox-3? Perception, 36.

Knapen, T., Gee, J. W. d., Brascamp, J., Nuiten, S., Hoppenbrouwers, S., & Theeuwes, J. (2016). Cognitive and ocular factors jointly determine pupil responses under equiluminance. PLOS ONE, 11(5), e0155574. 10.1371/journal.pone.0155574

Larson, E., Gramfort, A., Engemann, D. A., Leppakangas, J., Brodbeck, C., Jas, M., Brooks, T. L., Sassenhagen, J., McCloy, D., Luessi, M., King, J.-R., Höchenberger, R., Goj, R., Favelier, G., Brunner, C., van Vliet, M., Wronkiewicz, M., Rockhill, A., Holdgraf, C., … luzpaz. (2024). MNE-python (Version v1.8.0). Zenodo. 10.5281/zenodo.13340330

Liu, Y., Nour, M. M., Schuck, N. W., Behrens, T. E. J., & Dolan, R. J. (2022). Decoding cognition from spontaneous neural activity. Nature Reviews Neuroscience, 23(4), 204–214. 10.1038/s41583-022-00570-z

Maheu, M., Donner, T. H., & Wiegert, J. S. (2025). Serotonergic and noradrenergic interactions in pupil-linked arousal. 10.1101/2025.03.26.644382

Mante, V., Sussillo, D., Shenoy, K. V., & Newsome, W. T. (2013). Context-dependent computation by recurrent dynamics in prefrontal cortex. Nature, 503(7474), 78–84. 10.1038/nature12742

Maris, E., & Oostenveld, R. (2007). Nonparametric statistical testing of EEG- and MEG-data. Journal of Neuroscience Methods, 164(1), 177–190. 10.1016/j.jneumeth.2007.03.024

Matlab optimization toolbox. (2017). Natick, Massachusetts, United States, MathWorks.

Mejias, J. F., Murray, J. D., Kennedy, H., & Wang, X.-J. (2016). Feedforward and feedback frequency-dependent interactions in a large-scale laminar network of the primate cortex. Science Advances, 2(11), e1601335. 10.1126/sciadv.1601335

Miller, E. K., & Cohen, J. D. (2001). An integrative theory of prefrontal cortex function. Annual Review of Neuroscience, 24(1), 167–202. 10.1146/annurev.neuro.24.1.167

Mills, K. (2016). HCP-MMP1.0 projected on fsaverage. 10.6084/m9.figshare.3498446.v2

Minka, T. (2005). Divergence measures and message passing (MSR-TR-2005-173). https://www.microsoft.com/en-us/research/publication/divergence-measures-and-message-passing/

Mitra, P. P., & Pesaran, B. (1999). Analysis of dynamic brain imaging data. Biophysical Journal, 76(2), 691–708. 10.1016/S0006-3495(99)77236-X

Monov, G., Stein, H., Klock, L., Gallinat, J., Kühn, S., Lincoln, T., Krkovic, K., Murphy, P. R., & Donner, T. H. (2024). Linking cognitive integrity to working memory dynamics in the aging human brain. Journal of Neuroscience, 44(26). 10.1523/JNEUROSCI.1883-23.2024

Murphy, P. R., Krkovic, K., Monov, G., Kudlek, N., Lincoln, T., & Donner, T. H. (2024). Individual differences in belief updating and phasic arousal are related to psychosis proneness. Communications Psychology, 2(1), 1–14. 10.1038/s44271-024-00140-2

Murphy, P. R., Wilming, N., Hernandez-Bocanegra, D. C., Prat-Ortega, G., & Donner, T. H. (2021). Adaptive circuit dynamics across human cortex during evidence accumulation in changing environments. Nature Neuroscience, 1–11. 10.1038/s41593-021-00839-z

Nassar, M. R., Rumsey, K. M., Wilson, R. C., Parikh, K., Heasly, B., & Gold, J. I. (2012). Rational regulation of learning dynamics by pupil-linked arousal systems. Nature Neuroscience, 15(7), 1040–1046. 10.1038/nn.3130

Nienborg, H., & Cumming, B. G. (2009). Decision-related activity in sensory neurons reflects more than a neuron’s causal effect. Nature, 459(7243), 89–92. 10.1038/nature07821

O’Reilly, J. X., Schüffelgen, U., Cuell, S. F., Behrens, T. E. J., Mars, R. B., & Rushworth, M. F. S. (2013). Dissociable effects of surprise and model update in parietal and anterior cingulate cortex. Proceedings of the National Academy of Sciences, 110(38), E3660–E3669. 10.1073/pnas.1305373110

Ossmy, O., Moran, R., Pfeffer, T., Tsetsos, K., Usher, M., & Donner, T. H. (2013). The timescale of perceptual evidence integration can be adapted to the environment. Current Biology, 23(11), 981–986. 10.1016/j.cub.2013.04.039

Pedregosa, F., Varoquaux, G., Gramfort, A., Michel, V., Thirion, B., Grisel, O., Blondel, M., Prettenhofer, P., Weiss, R., Dubourg, V., Vanderplas, J., Passos, A., Cournapeau, D., Brucher, M., Perrot, M., & Duchesnay, É. (2011). Scikit-learn: Machine learning in python. Journal of Machine Learning Research, 12(85), 2825–2830. Retrieved 2024, from http://jmlr.org/papers/v12/pedregosa11a.html

Peixoto, D., Verhein, J. R., Kiani, R., Kao, J. C., Nuyujukian, P., Chandrasekaran, C., Brown, J., Fong, S., Ryu, S. I., Shenoy, K. V., & Newsome, W. T. (2021). Decoding and perturbing decision states in real time. Nature, 591(7851), 604–609. 10.1038/s41586-020-03181-9

Pelli, D. G. (1997). The VideoToolbox software for visual psychophysics: Transforming numbers into movies. Spatial Vision, 10(4), 437–442. 10.1163/156856897X00366

Pfeffer, T., Ponce-Alvarez, A., Tsetsos, K., Meindertsma, T., Gahnström, C. J., van den Brink, R. L., Nolte, G., Engel, A. K., Deco, G., & Donner, T. H. (2021). Circuit mechanisms for the chemical modulation of cortex-wide network interactions and behavioral variability. Science Advances, 7 (29), eabf5620. 10.1126/sciadv.abf5620

Pfurtscheller, G., & Lopes da Silva, F. H. (1999). Event-related EEG/MEG synchronization and desynchronization: Basic principles. Clinical Neurophysiology, 110(11), 1842–1857. 10.1016/S1388-2457(99)00141-8

Purcell, B. A., & Kiani, R. (2016). Hierarchical decision processes that operate over distinct timescales underlie choice and changes in strategy. Proceedings of the National Academy of Sciences, 201524685. 10.1073/pnas.1524685113

Rao, R. P. N., & Ballard, D. H. (1999). Predictive coding in the visual cortex: A functional interpretation of some extra-classical receptive-field effects. Nature Neuroscience, 2(1), 79–87. 10.1038/4580

Ratcliff, R., & McKoon, G. (2008). The diffusion decision model: Theory and data for two-choice decision tasks. Neural Computation, 20(4), 873–922. 10.1162/neco.2008.12-06-420

Resulaj, A., Kiani, R., Wolpert, D. M., & Shadlen, M. N. (2009). Changes of mind in decision-making. Nature, 461(7261), 263–266. 10.1038/nature08275

Sakai, K. (2008). Task set and prefrontal cortex. Annual Review of Neuroscience, 31, 219–245. 10.1146/annurev.neuro.31.060407.125642

Sara, S. J., & Bouret, S. (2012). Orienting and reorienting: The locus coeruleus mediates cognition through arousal. Neuron, 76(1), 130–141. 10.1016/j.neuron.2012.09.011

Sarafyazd, M., & Jazayeri, M. (2019). Hierarchical reasoning by neural circuits in the frontal cortex. Science, 364(6441), eaav8911. 10.1126/science.aav8911

Schwabe, L., Haddad, L., & Schachinger, H. (2008). HPA axis activation by a socially evaluated cold-pressor test. Psychoneuroendocrinology, 33(6), 890–895. 10.1016/j.psyneuen.2008.03.001

Ségonne, F., Dale, A. M., Busa, E., Glessner, M., Salat, D., Hahn, H. K., & Fischl, B. (2004). A hybrid approach to the skull stripping problem in MRI. NeuroImage, 22(3), 1060–1075. 10.1016/j.neuroimage.2004.03.032

Shadlen, M. N., & Kiani, R. (2013). Decision making as a window on cognition. Neuron, 80(3), 791–806. 10.1016/j.neuron.2013.10.047

Shadlen, M. N., & Newsome, W. T. (2001). Neural basis of a perceptual decision in the parietal cortex (area LIP) of the rhesus monkey. Journal of Neurophysiology, 86(4), 1916–1936. 10.1152/jn.2001.86.4.1916

Silver, M. A., & Kastner, S. (2009). Topographic maps in human frontal and parietal cortex. Trends in Cognitive Sciences, 13(11), 488–495. 10.1016/j.tics.2009.08.005

Smith, S. M., & Nichols, T. E. (2009). Threshold-free cluster enhancement: Addressing problems of smoothing, threshold dependence and localisation in cluster inference. NeuroImage, 44(1), 83–98. 10.1016/j.neuroimage.2008.03.061

Steinemann, N., Stine, G. M., Trautmann, E., Zylberberg, A., Wolpert, D. M., & Shadlen, M. N. (2024). Direct observation of the neural computations underlying a single decision. eLife, 12, RP90859. 10.7554/eLife.90859

Stolk, A., Todorovic, A., Schoffelen, J.-M., & Oostenveld, R. (2013). Online and offline tools for head movement compensation in MEG. NeuroImage, 68, 39–48. 10.1016/j.neuroimage.2012.11.047

Tavoni, G., Doi, T., Pizzica, C., Balasubramanian, V., & Gold, J. I. (2022). Human inference reflects a normative balance of complexity and accuracy. Nature Human Behaviour, 1–16. 10.1038/s41562-022-01357-z

Urai, A. E., & Donner, T. H. (2022). Persistent activity in human parietal cortex mediates perceptual choice repetition bias. Nature Communications, 13(1), 6015. 10.1038/s41467-022-33237-5

Usher, M., & McClelland, J. L. (2001). The time course of perceptual choice: The leaky, competing accumulator model. Psychological Review, 108. 10.1037/0033-295X.108.3.550

van den Brink, R. L., Hagena, K., Wilming, N., Murphy, P. R., Büchel, C., & Donner, T. H. (2023). Flexible sensory-motor mapping rules manifest in correlated variability of stimulus and action codes across the brain. Neuron, 111(4), 571–584.e9. 10.1016/j.neuron.2022.11.009

van den Brink, R. L., Murphy, P. R., & Nieuwenhuis, S. (2016). Pupil diameter tracks lapses of attention. PLOS ONE, 11(10), e0165274. 10.1371/journal.pone.0165274

Van Essen, D. C., Ugurbil, K., Auerbach, E., Barch, D., Behrens, T. E. J., Bucholz, R., Chang, A., Chen, L., Corbetta, M., Curtiss, S. W., Della Penna, S., Feinberg, D., Glasser, M. F., Harel, N., Heath, A. C., Larson-Prior, L., Marcus, D., Michalareas, G., Moeller, S., … Yacoub, E. (2012). The human connectome project: A data acquisition perspective. NeuroImage, 62(4), 2222–2231. 10.1016/j.neuroimage.2012.02.018

Van Veen, B. D., van Drongelen, W., Yuchtman, M., & Suzuki, A. (1997). Localization of brain electrical activity via linearly constrained minimum variance spatial filtering. IEEE transactions on bio-medical engineering, 44(9), 867–880. 10.1109/10.623056

van Kempen, J., Gieselmann, M. A., Boyd, M., Steinmetz, N. A., Moore, T., Engel, T. A., & Thiele, A. (2021). Top-down coordination of local cortical state during selective attention. Neuron, 109(5), 894–904.e8. 10.1016/j.neuron.2020.12.013

Vyas, S., Golub, M. D., Sussillo, D., & Shenoy, K. V. (2020). Computation through neural population dynamics. Annual Review of Neuroscience, 43, 249–275. 10.1146/annurev-neuro-092619-094115

Wandell, B. A., Dumoulin, S. O., & Brewer, A. A. (2007). Visual field maps in human cortex. Neuron, 56(2), 366–383. 10.1016/j.neuron.2007.10.012

Wang, X.-J. (2022). Theory of the multiregional neocortex: Large-scale neural dynamics and distributed cognition. Annual Review of Neuroscience, 45(1), 533–560. 10.1146/annurev-neuro-110920-035434

Weiss, A., Chambon, V., Lee, J. K., Drugowitsch, J., & Wyart, V. (2021). Interacting with volatile environments stabilizes hidden-state inference and its brain signatures. Nature Communications, 12(1), 2228. 10.1038/s41467-021-22396-6

Wilming, N., Murphy, P. R., Meyniel, F., & Donner, T. H. (2020). Large-scale dynamics of perceptual decision information across human cortex. Nature Communications, 11(1), 5109. 10.1038/s41467-020-18826-6

Wyart, V., de Gardelle, V., Scholl, J., & Summerfield, C. (2012). Rhythmic fluctuations in evidence accumulation during decision making in the human brain. Neuron, 76(4), 847–858. 10.1016/j.neuron.2012.09.015

Xia, R., Chen, X., Engel, T. A., & Moore, T. (2024). Common and distinct neural mechanisms of attention. Trends in Cognitive Sciences, 28(6), 554–567. 10.1016/j.tics.2024.01.005

Yu, A. J., & Dayan, P. (2005). Uncertainty, neuromodulation, and attention. Neuron, 46(4), 681–692. 10.1016/j.neuron.2005.04.026

Zhang, J., Kriegeskorte, N., Carlin, J. D., & Rowe, J. B. (2013). Choosing the rules: Distinct and overlapping frontoparietal representations of task rules for perceptual decisions. Journal of Neuroscience, 33(29), 11852–11862. 10.1523/JNEUROSCI.5193-12.2013

